# Early life nociception is influenced by peripheral growth hormone signaling

**DOI:** 10.1101/2020.07.16.206904

**Authors:** Adam J. Dourson, Zachary K. Ford, Kathryn J. Green, Carolyn E. McCrossan, Megan C. Hofmann, Renita C. Hudgins, Michael P. Jankowski

**Affiliations:** Department of Anesthesia; Center for Understanding Pediatric Pain, Cincinnati Children’s Hospital Medical Center; Department of Pediatrics, University of Cincinnati, College of Medicine, Cincinnati, OH 45229

## Abstract

A number of cellular systems work in concert to modulate nociceptive processing in the periphery, but the mechanisms that regulate neonatal nociception may be distinct compared to adults. Our previous work indicated a relationship between neonatal hypersensitivity and growth hormone (GH) signaling. Here, we explored the peripheral mechanisms by which GH modulated neonatal nociception under normal and injury conditions (incision). We found that GH receptor signaling in primary afferents maintains a tonic inhibition of peripheral hypersensitivity. After injury, a macrophage dependent displacement of injury-site GH was found to modulate neuronal transcription at least in part via serum response factor regulation. A single GH injection into the injured hindpaw muscle effectively restored available GH signaling to neurons and prevented acute pain-like behaviors, primary afferent sensitization, and neuronal gene expression changes. GH treatment also inhibited long-term somatosensory changes observed after repeated peripheral insult. Results may indicate a novel mechanism of neonatal nociception.

**Significance statement:** Although it is noted that mechanisms of pain development in early life are unique compared to adults, little research focuses on neonatal-specific peripheral mechanisms of nociception. This gap is evident in the lack of specialized care for infants following an injury including surgeries. This report evaluates how distinct cellular systems in the periphery including the endocrine, immune and nervous systems work together to modulate neonatal-specific nociception. We uncovered a novel mechanism by which muscle injury induces a macrophage-dependent sequestration of peripheral growth hormone that effectively removes its normal tonic inhibition of neonatal nociceptors to promote acute pain-like behaviors. Results indicate a possible new strategy for treatment of neonatal post-surgical pain.

## Introduction

Functional restoration after injury requires a coordinated response between immune cells, neurons, and local tissues within the affected area. This response further generates a nociceptive signal via primary sensory neurons that is required to inform the organism of the ongoing repair process (Baoge et al 2012) (Sass et al 2018) (McMahon et al 2015) (Basbaum et al 2009) (Philippou et al 2012). Responses from each of these individual cell types play a role in how noxious signals are transduced into the central nervous system (CNS) ^e.g.^ (Basbaum et al 2009).

The immature dorsal root ganglion (DRG) contains a compilation of functional sensory neuron subtypes that is distinct from adults (Jankowski et al 2014) (Sharma et al 2020). As such, neonates are particularly vulnerable to sensory impairment during developmental injury (Lim & Godambe 2017). Recent work in animals (Ren et al 2004) (Walker et al 2009) and humans (Hermann et al 2006) (Walker et al 2018) (Moriarty et al 2018) (Walker 2019) indicates that early life injury enhances pain-related responses later in life. These “priming” effects have been linked to alterations in the central nervous system (Walker et al 2016) (Baccei 2016) (Brewer & Baccei 2019) but the peripheral component is less studied (Moriarty et al 2018). We have shown that the pattern of primary afferent sensitization after neonatal injury is unique to that observed in adult nociceptors (Jankowski et al 2014) (Ye & Woodbury 2010) (Koerber et al 2010) (Koerber & Woodbury 2002). This suggests that the mechanisms of nociception in neonates may be somewhat distinct.

We recently found that growth hormone (GH) may be one factor involved in generalized pain-related responses to peripheral injury in neonates (Liu et al 2017) (Ford et al 2019). The largest increases in systemic GH levels are known to occur during early postnatal development, which corresponds with the most rapid growth period (Bartholomew EF 2009). This is the same developmental period when GH was found to substantially influence pain-like responses and primary afferent function (Liu et al 2017) (Ford et al 2019). GH receptors (GHr) typically affect cellular functions through activation of various transcription factors such as the signal transducers and activators of transcription (STATs), serum response factor (SRF) or ERK like kinases (ELKs) (Ceseña et al 2007). These factors can also be invoked in response to an aversive stimulus and can modulate both transcriptional and behavioral phenotypes (Salaffi et al 2018) (Gomez et al 2018). Immune cells that infiltrate injury sites release various cytokines and growth factors during repair that can themselves be pro-nociceptive (Ren & Dubner 2010) (McMahon et al 2005) (McMahon et al 2015) (Basbaum et al 2009) (Philippou et al 2012). Interestingly, macrophages in particular are known to use peripheral GH to modulate local inflammation (Govers et al 1999) (Strous et al 1996) (Lu et al 2013) (Schneider et al 2019). Together, altered GH signaling within primary afferent neurons may modulate peripheral sensitization in coordination with the immune system (Basbaum et al 2009) (McMahon et al 2015) (Ford et al 2019).

Clinical reports show that in addition to growth problems, many children with GH deficiency (Pinho-Ribeiro et al 2018) (Cuatrecasas 2009) (Dattani & Preece 2004) report pain (Cimaz et al 2001). Other studies have found that exogenous GH treatment may be an effective pain therapy for patients with erythromelalgia (Cimaz et al 2001), fibromyalgia (Cuatrecasas et al 2014) (Cuatrecasas et al 2010) (Cuatrecasas et al 2012) (Cuatrecasas et al 2007) or low back pain (Dubick et al 2015). Conversely, GH receptor blockers, used to treat acromegaly, can produce pain ^e.g.^ (van der Lely et al 2001). Thus, understanding the mechanisms of how GH regulates pain-like responses may help us understand basic nociceptive processing in the immature nervous system. Here, we tested the hypothesis that GH modulates peripheral sensitization after neonatal muscle incision through unique endocrine-peripheral nervous-and immune system interactions.

## Materials and Methods

### Animals

Male and female mice that were postnatal day 7 (P7) to P14 (+/− 1d around the specified age range) or P35-P56 were used throughout all experiments. In all cases except for experiments regarding SRF knockdown, no sex differences were observed and thus data is combined from both males and females for ease of presentation. Neonatal animals were kept with the dam and only separated for short durations at a time (<1.5hrs) to perform behavioral experiments. All mice were kept in an environment-controlled facility at Cincinnati Children’s Hospital Medical Center with free access to food and water while on a 12-hour light/dark cycle. Animals are defined as wildtype (WT) controls, heterozygous (+/−), or homozygous (−/−) for the genetic manipulation throughout all experiments.

Swiss Webster mice were born in house or purchased from Charles River (Wilmington, MA) or Envigo (Indianapolis, IN) and were used as an outbred strain for most experimentation. Littermate and non-littermate C57BL/6 animals born in-house were used as controls for genetic lines bred on that background. A deletion of the growth hormone receptor (GHr) specifically on macrophages was kindly gifted to use by Dr. Ram Menon (University of Michigan). This mouse was generated by crossing a LysM-Cre positive animal (see Jax Stock#: 004781) with a GHr floxed animal to induce cell type specific genetic deletion of GHr in monocytes/macrophages. Animals with a knockout of the growth hormone releasing hormone receptor (GHRHr) were purchased from The Jackson Laboratory (C57BL/6J-Ghrhr^lit^/J; Stock#: 000533). We developed a sensory neuron specific deletion of GHr by crossing a tamoxifen inducible Cre recombinase driven by the Advillin (Adv) promotor purchased from The Jackson Laboratory (Advillin-CreERT2; Stock#: 026516). Mice were crossed in-house with cryo-recovered (CCHMC Transgenic Core Facility) GHr floxed (GHr^tm1b(KOMP)Wtsi^) embryos purchased from the KOMP Repository (Design ID#: 49728) to make an inducible sensory neuron specific GHr knockout (Adv;GHr^f/f^). A Cre dependent reporter mouse was also used to drive tdTomato (tdTOM) expression (B6.Cg-Gt(ROSA)26Sor^tm14(CAG-tdTomato)Hze^/J) and was purchased from The Jackson Laboratory (Stock#: 007914). Finally, LysM-Cre animals purchased from The Jackson Laboratory (Stock#: 004781) were crossed to the tdTom mice to generate myeloid/macrophage reporter mice. All procedures were approved by the CCHMC Institutional Animal Care and Use Committee in compliance with AALAC approved practices.

### Behavioral Measures

Neonatal animals (P7-P14) were transferred from their home cage to opaque chambers with a translucent lid and acclimated in a temperature-controlled environment for 10 minutes prior to assessments. Adolescent animals (≥P35) were transferred to raised translucent boxes with a grid mesh bottom and acclimated for 25 minutes prior to data collection. After habituation, behavioral experiments followed including spontaneous paw guarding assessment, muscle mechanical withdrawal thresholds, cutaneous mechanical withdrawal thresholds, grip strength and/or proprioceptive behaviors. No cohort received more than two of the listed behaviors at a time to reduce stress and maternal separation time for neonatal animals. Data was obtained at baseline, 1d and/or 3d, or up to 21d for adolescent animals, post injury as indicated.

Spontaneous paw guarding assessments scores preferential weight bearing on a scale of zero to two, where zero is no guarding after injury, 1 indicates shifted weight bearing but the paw still touches the floor and 2 indicates full paw lifting. Assessments were made for a duration of 1 minute. Neonatal animals were scored every five minutes for 30 minutes, and adolescent animals were scored for an hour.

Hind paw muscle withdrawal was assessed with a digital Randall Selitto device (IITC Life Science Inc. Woodland Hills, CA, USA) with a dulled probe attachment ~2 mm wide at the tip. The dorsal paw was supported by the upper machine arm and the medial plantar paw was slowly pressed with the dulled probe until a robust withdrawal response was evoked. The gram force that elicited a withdrawal response was considered threshold. Three trials were obtained at least five minutes intervals and averaged together for analysis. Maximum squeezing force was 150g for neonates and 350g or 500g for adolescent mice of different strains.

Cutaneous mechanical withdrawal thresholds were assessed on the dorsal surface of the hindpaw as described in previous reports (Liu et al 2017) (Jankowski et al 2014) (Marsh et al 1999) for neonatal animals using an increasing series of calibrated von Frey filaments ranging from 0.07g to 6g. Threshold to withdrawal was determined in three trials with five minutes intervals between trials and averaged.

Dynamic paw muscle strength was assessed by neonatal hanging time (Au - Feather-Schussler & Au - Ferguson 2016). Animals were held with forepaws near a thin metal rod spanning a 9.5 cm diameter apparatus until the rod was gripped. Animals were timed while freely hanging above a 12 cm padded drop until they released the bar, escaped the apparatus by climbing out, or 60 seconds was reached. An escape was determined to require enough muscle strength to pull up and out of the apparatus and was thus set to maximum time. Three trials were recorded and averaged.

To evaluate proprioception, we recorded the animals’ innate righting reflex (Dallman & Ladle 2013). Animals were gently turned over and placed on their back. The time to flip over to all four limbs was recorded and averaged over three trials with 5-minute intervals between trials.

Animals were weighed and their temperature was taken after an injection of growth hormone to determine if any GH-related side effects were present with our injection strategy. 24 hours after an intramuscular hindpaw injection, animals were weighed on a tabletop scale and their temperature was recorded using a digital surface contact thermometer pressed against the chest of the anesthetized animal according to previous methods (Liu et al 2017) (Goodrich 1977).

### Injections

Growth hormone was injected directly into the hind paw muscles in uninjured mice or in animals receiving incision injury. Experiments in Fig. 5c were intraperitoneal GH injections. Dosing ranged from 0.1mg/kg to 1.5 mg/kg in 10 μL for all neonatal experiments. Adolescent animals were injected with the 1.5mg/kg dose in 18 μL. 5mg/kg GH binding protein (GHBP) or vehicle (0.1% bovine serum albumin (BSA)) in PBS was injected in 10 μL into the hindpaw muscle. To induce Cre-recombinase in AdvGHr animals, tamoxifen was made fresh at 25mg/mL in corn oil and uninjured animals were singly injected intraperitoneal (i.p.) at P7 at a dose of 250mg/kg tamoxifen (Hester & Danzer 2013).

### Surgical Hind Paw Incisions

Animals were anesthetized with 2-3% isoflurane and a longitudinal incision of the right hairy hind paw skin was made lateral to the main saphenous nerve innervation territory. Then incision was continued in between the bones through to the flexor digitorum brevis muscles. Blunt manipulation of the muscle was performed using #5 forceps, but the plantar skin was left untouched. Prior to wound closing with 7-0 sutures, interventions corresponding to the experiment were injected into the incision site. When appropriate, adolescent surgical hind paw incisions were performed using the same procedures, with wounds closed with 6-0 sutures. Animals were allowed to recover for the indicated times. For comparisons, some cohorts only received the hairy skin incision or a single suture through intact skin (sham) but did not experience the muscle incision. For dual incision assays, similar procedures were followed as described above except the first incision was made at P7 and the second incision (when indicated) was performed at P35.

### Sciatic Nerve Injections

Mice were placed on a warming pad and kept under 2-3% isoflurane anesthesia as the right sciatic nerve was revealed by a small incision of the skin and cautious separation of the underlying muscle. Carefully, as to not stretch the nerve, the sciatic was separated from surrounding tissue and raised onto a malleable plastic platform. Targeting or control siRNAs were injected directly into the sciatic nerve above the trifurcation using quartz microelectrodes connected to a picospritzer with 8-10 short pulses at 1-2psi. Approximate volume of injection was ~100nL. 4 different duplexes from ON-TARGETplus SMARTpool siRNAs (Dharmacon, Lafayette; Catalog#: 4390771) against serum response factor were first tested *in vitro* (Neuro2a cells) to determine knock down efficiency of the individual siRNA duplexes similar to previous reports ^e.g.^ (Queme et al 2016) (Ross et al 2016) (Liu et al 2017) (data not shown). The most efficient sequence was determined using realtime PCR and used for all subsequent *in vivo* analyses (sense: 5’-S-S-GCAGCAACCUCACCGAGCUUU; antisense: 5’-P-AGCUCGGUGAGGUUGCUGCUU). siRNAs were first conjugated to Penetratin-1 according to manufacturer’s instructions after thiol removal (Dharmacon) and reconstituted at 90μM. Depending on age, siRNAs were then injected into the sciatic nerve as described above one (<P10) or two (≥P10) days prior to incision and further experimentation to allow for retrograde transport of the siRNAs to the DRG somas. The non-coding control siRNA has been used previously and does not target any murine gene (ThermoFisher D-001206-14-05).

### Realtime RT-PCR

RNA was isolated from lumbar 3/4/5 (L3/4/5) dorsal root ganglia (DRG) on the side ipsilateral to injury. RNeasy Mini Kit (Qiagen Stock#: 74104) was performed on DRGs for total mRNA isolation and RNeasy Fibrous Tissue Mini Kit (Qiagen Stock#: 74704) was used to isolate muscle mRNA. All RNA isolations were performed exactly according to the manufactures directions. For standard realtime PCR assessments, 500ng of total RNA was reverse transcribed into cDNA and realtime PCR performed using SYBR Green Master Mix on a StepOne realtime PCR system (Applied Biosystems). Quantitative PCR was analyzed by the ΔΔcycle threshold (CT) method with normalization to GAPDH. Fold change between conditions was determined and converted to a percent change where 2-fold = 100% change. Primer sequences are all recorded in Extended Data Table 1-1.

### Western Blot

Flexor digitorum brevis muscle or L3/4/5 DRGs were dissected and frozen on dry ice. After homogenization in protein lysis buffer as completed previously ^e.g.^ (Ross et al 2018), 20 μg samples were boiled in gel loading buffer containing β-mercaptoethanol as a reducing agent and loaded onto a 12% or “AnyKD” precast polyacrylamide gel (Bio-Rad 4569033) for western blot analysis. Gels were transferred to a polyvinylidene difluoride membranes (PVDF; Merck Millipore Ltd., Tullagreen, Ireland) at 35V for 16-18 hours at 4°C. The next day, transfer quality was assessed by staining (Coomassie Brilliant Blue BioRad 1610436) the gel for any remaining proteins. The membrane was washed, blocked with Odyssey Blocking Buffer (BB) (LiCor 927-40000) diluted in PBS (1:4), and incubated in 2x PBS with 0.2%Tween and BB (1:1) with primary antibodies. After incubation overnight, the membranes were washed, incubated in 2x PBS with 0.2%Tween and 0.01% SDS and BB (1:4) with appropriate infrared-conjugated secondary antibodies (LiCor) and visualized on LiCor Odyssey CLx protein imaging system. Exposure times were consistent between runs and gain was always set to 1.0. Band intensity was then quantified using ImageJ software (NIH) similar to previous procedures (Liu et al 2017). Primary antibodies included GH (LS‑C146263, polyclonal rabbit, 1:1,000) and GAPDH (Abcam 83956, polyclonal chicken, 1:2,000).

### Immunohistochemistry

DRGs, sciatic nerve and hindpaw muscle were sectioned on a cryostat at 10 μm (DRG and sciatic nerve) or 20 μm (muscle). DRGs were frozen on dry ice in OCT medium and muscle was snap frozen in liquid nitrogen. All tissue was kept at −80°C until use. Cryostat sections were then fixed on the slide. Slides were washed and blocked prior to overnight primary antibody incubation. The next day, the tissue was washed and stained for secondary antibodies before cover slipping with mounting media containing DAPI to mark nuclei (Fisher Scientific 17985-50). For immunocytochemistry, samples were processed by Cincinnati Children’s Pathological core. Briefly, slides were pretreated with citrate buffer, washed and incubated with primary antibody for 32 minutes. Detection was completed with the DAB rabbit kit (Ventana#: 760-151), counterstained with hematoxylin and blued with bluing reagent, and then dehydrated before coverslipping. Primary antibodies used were: GH (LS‑C146263, polyclonal rabbit, 1:500 (fluorescence) or 1:100 (DAB)), GHr (Abcam 202964, polyclonal rabbit, 1:1,000), SRF (Abcam 53147, polyclonal rabbit, 1:250), and dystrophin (Abcam 15277, polyclonal rabbit, 1:250). Fluorescent imaging was observed on a Nikon confocal microscope and all gain and laser power were maintained equally across all samples in each experiment. For quantification, images were converted to grayscale by a blind investigator and an equal threshold was applied to all images. A 200×200 region of interest (ROI) defined by positive staining was analyzed for particles of sufficient size and were quantified for mean gray value (ImageJ User Guide).

To quantify myofiber cross sectional area, we used the protocol established by Nikolaou and colleagues (Nikolaou et al 2015). Briefly, dystrophin stained sections of hindpaw muscle were acquired on the Nikon confocal microscope at high intensity to obtain consistent signal around myofibers. Images were captured using NIS Elements software and then using Fiji software (Schindelin et al 2012), images were converted to 8-bit binary images. Converted images were then manually edited to remove non-muscle regions or damaged fiber staining. Myofiber cross-sectional area measurements were then obtained for each fiber in the section. Three non-consecutive sections per condition were analyzed and averaged.

### *Ex vivo* Preparation

A novel neonatal *ex vivo* hind paw muscle-tibial nerve-DRG-spinal cord recording preparation was used to directly assess the response properties of individual primary afferent neurons under our various conditions. Briefly, based on the forepaw prep previously described ^e.g.^(Queme et al 2020), animals were first anesthetized with a mix of ketamine and xylazine (100 and 16 mg/kg, respectively) and then perfused with ice cold oxygenated (95%o_2_/5% Co_2_) artificial cerebral spinal fluid (aCSF; in mM: 127.0 NaCl, 1.9 KCl, 1.2 KH2PO4, 1.3 MgSO4, 2.4 CaCl2, 26.0 NaHCO3, and 10.0 D-glucose). The intact spine and right hind leg were isolated and transferred to a new dish with circulating oxygenated aCSF. The hind paw muscle (with bone intact), tibial/sciatic nerve, L1-L6 DRGs and corresponding spinal cord segments were dissected in continuity. The spinal cord was hemisected, and the intact preparation was then transferred to a new recording chamber under the same conditions. The paw with revealed muscle was pinned to a metal grate within an inner bath under its own circulation of o_2_aCSF. The nerve was fed through a small gap of the inner bath and the spinal cord and DRGs were pinned within the outer dish. The hole between the dishes was filled with petroleum jelly to separate the baths and hold the nerve in place. The bath was slowly warmed to 32°C.

Quartz microelectrodes (impedance>150MΩ) containing 5% Neurobiotin (Vector Laboratories, Burlingame, CA) in 1M potassium acetate were used for sharp electrode single unit recordings in the L3 or L4 DRGs. An impaled cell body was determined to have axons in the tibial nerve by a response from electrical search stimulus by a suction electrode placed on the side of the nerve. Once a cell was determined to have axons in the tibial nerve, the muscle was probed with concentric bipolar electrode to locate the cell’s receptive field (RF) in the hindpaw muscle. Then the muscle was probed with mechanical stimuli, thermal stimuli and chemical stimuli in this order. For mechanical stimuli, an increasing series of von Frey filaments ranging from 0.07-10 grams were used to stimulate the RF for ~1-2s. Then cold (~2°C) followed by hot (~53°C) physiological saline was delivered to the RF. Following the thermal stimulations, two distinct metabolite mixtures were slowly introduced into the inner bath surrounding the hind paw muscles. First a “low” concentration of metabolites (15 mM lactate, 1 μM ATP, pH 7.0) were applied for approximately two minutes and then washed out. After washout, a “high” concentration of metabolites (50 mM lactate, 5 μM ATP, pH 6.6) was added to the inner chamber in the same manner. Metabolites were oxygenated and heated to physiological conditions with an in-line heater to maintain bath conditions. ATP was added just before perfusion of the muscle. After metabolite stimulation, mechanical and thermal responsiveness was again assessed.

All activity was recorded by Spike2 software (Cambridge Electronic Design) and was later analyzed offline. Response latencies were recorded and divided by nerve length to determine conductance velocities to categorize group IV afferents (≤1.2 m/s) or group III afferents (1.2-14 m/s). Mechanical thresholds were determined by the least amount of force necessary to elicit at least two action potentials. Peak instantaneous frequencies (IF) were determined to assess the maximum response to a peripheral stimulus while firing rates (FR) were determined to obtain the maximum number of events that occurred over a given period of time (200ms bins). The distribution analyses were determined to be the number of cells that responded to a given stimuli divided by the number of total cells receiving that stimulus. Since chemical stimulation was not given to every cell tested, the distribution of each subtype of chemically activated cells (low, high or both) was divided by total chemically responsive cells per condition. No differences in response properties were found between cells obtained at the beginning of a recording experiment compared to the end of the session. Based on previous work ^e.g.^(Queme et al 2020) (Jankowski et al 2014) and power analyses, we determined that ≥50 cells per group would be required.

### Experimental Design and Statistical Analysis

Data were analyzed using SigmaPlot software (v.14). Critical significance was set to α<0.05. All data was first checked for normal distribution with Shapiro-Wilk and equal variance with Brown-Forsythe and then parametric or nonparametric tests were used accordingly. Specific tests are indicated in the figure legends. For behavioral data containing the same animals treated with an intervention over time, a two-way repeated measures analysis of variance (RM ANOVA) was used. In behavioral data in which only one time point or a percent change from baseline/naïve was compared across groups a one-way ANOVA was used. Analysis of cDNA, protein quantification, area under the curve, or *ex vivo* analysis was measured by a one-way ANOVA or corresponding non-parametric test across groups. All analyses that passed the omnibus F-test were further discriminated by Tukey’s, Holm-Sidak, or Dunn’s post hoc analysis as noted in figure legends. Discrete categorical data were analyzed by chi-square across all groups. Graphical panels were made using Graphpad Prism (v.8) and compiled in Adobe Photoshop. In all studies involving subjective measures including all behavior, *ex vivo* and IHC quantification, the researcher was blinded by co-investigators or by the unknown genotype of the animal. The rare occasion of a detected outlier defined as being greater than 2 standard deviations away from the mean was removed. Biological replicates (“n”) are provided for each figure in the legends or text. Technical replicates are explained above for each experiment.

## Results

### Sensory neuron specific deletion of growth hormone receptor modulates peripheral responsiveness in uninjured neonatal mice

Previously, we have demonstrated that a systemic reduction in growth hormone levels resulted in neonatal hypersensitivity (postnatal day 7 (P7)-P14) to peripheral stimuli that resolved by P21 (Ford et al 2019).To determine if the effects of reduced GH-signaling on nociception were peripherally mediated, we injected growth hormone binding protein (GHBP) locally into the right hind paw of uninjured neonatal (P7-P14) mice: the time frame in which we observed GH deficiency-related hypersensitivity. We found that one day after injection, GHBP resulted in spontaneous paw guarding behaviors (Fig. 1a). To then test if the anti-nociceptive effects of GH were due to a direct effect on primary afferents, we developed a transgenic mouse that allowed for targeted deletion of GHr in a time dependent manner in sensory neurons. We crossed the advillin (Adv)-creERT2 mouse (Lau et al 2011) with a floxed GHr line and injected neonatal pups with tamoxifen at P7 to initiate sensory neuron specific deletion of GHr (Fig. 1b). Using this strategy, we first confirmed that control animals expressed normal levels of GHr in the DRGs while tamoxifen injected Adv;GHr^f/f^ neonates had significantly reduced GHr expression as assessed with immunocytochemistry (Fig. 1b’,b’’). Realtime PCR analysis confirmed that within 5-7 days post tamoxifen injection, that GHr mRNA was significantly reduced in the DRGs compared to controls (−45 ± 25% vs. controls; n=5-13 per group; p<0.05). This corresponded with an upregulation of insulin-like growth factor receptor 1 (IGFr1) (136 ± 46% vs. control; p<0.05) as well as serum response factor (SRF) mRNA (229 ± 52% vs. control; p<0.05) confirming previous literature (Carter-Su et al 2016) that these factors are downstream of growth hormone signaling. Interestingly, STAT5 was not altered in the DRGs of tamoxifen injected Adv;GHr^f/f^ mice (7 ± 41% vs. control; p>0.05). We then performed behavioral analyses in these animals and found that although proprioception (righting reflexes) was unaffected by the sensory neuron targeted knockout of GHr (Fig. 1c), we detected alterations in other behaviors. Static cutaneous mechanical responsiveness (von Frey filament withdrawal thresholds), and withdrawal thresholds to probing of the muscles (muscle squeezing) were significantly altered by GHr knockout in uninjured animals (Fig. 1d,e). Further, in a dynamic neonatal muscle strength assay, we again found that, unlike littermate controls, animals with sensory neuron specific GHr knockout did not significantly increase in strength over time (Fig. 1f). Area under the curve (AUC) analysis indicates that neonates with GHr mutations have lower overall cutaneous and muscle withdrawal thresholds and muscle strength scores compared to control animals (Fig. 1d’,e’,f’).

**Figure 1:**
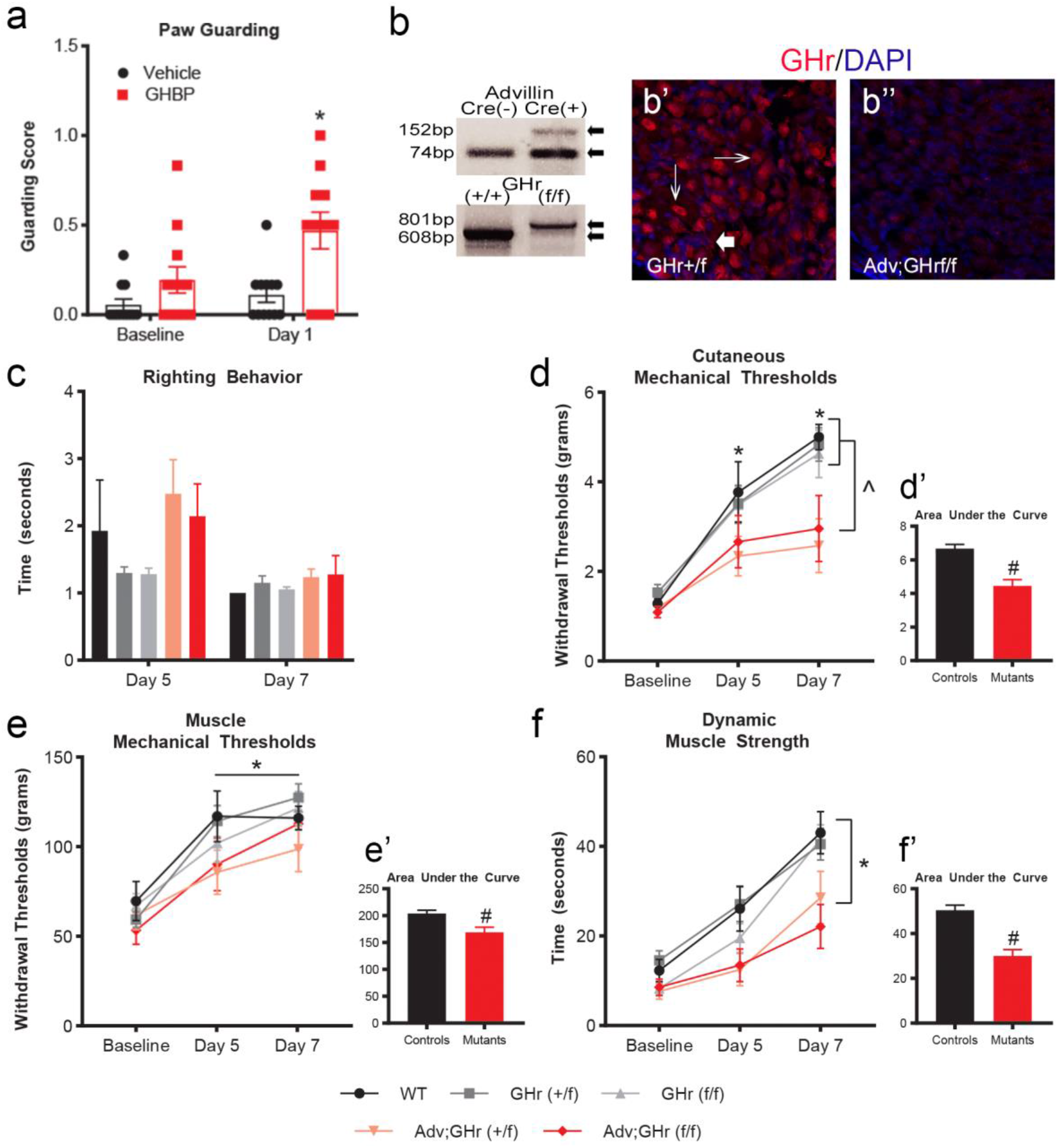
Modulation of neonatal nociception by peripheral GH signaling. a, Animals injected with GHBP display increased paw guarding at 1d compared to BL while vehicle injected animals do not and guard significantly more than vehicle injected groups. * indicates p<0.001 vs. BL and p<0.05 vs. Day 1 control. n = 12/group, Two-way RM ANOVA, Tukey’s post hoc. b, Representative genotyping analysis of Adv-Cre; GHr^f/f^ mice. b’,b’’, Immunostaining of DRGs for GHr (red) and nuclear labeling using DAPI (blue) in tamoxifen treated control (Adv;GHr^+/f^) and Adv;GHr^f/f^ mice. Arrows indicate GHr+ neuronal staining, large arrow indicates nerve fiber. c, Righting reflexes are not different across groups at 5- and 7-days post tamoxifen. n = 11-14 (GHr+/+), 26-30 (GHrf/+), 26-29 (GHrf/f), 13-16 (Adv;GHrf/+), and 5-7 (Adv;GHrf/f)/time point, Two-way RM ANOVA, Tukey’s post hoc. d, Cutaneous mechanical withdrawal thresholds increased over time in control groups but not in Adv;GHr^f/+^ (n = 8-12/time point) nor Adv;GHr^f/f^ (8-10/time point) animals. *p<0.001 vs. BL; ^p<0.05 vs. each control group. Two-way RM ANOVA, Tukey’s post hoc. d’, Combined mutant (Adv;GHr^f/+^, Adv;GHr^f/f^) groups have significantly lower cutaneous mechanical withdrawal thresholds compared to control groups (GHr^+/+^, GHr^f/+^, GHr^f/f^, n=6-7, 16-22, 10-14/time point respectively) in an area under the curve from BL (P7) to 7d (P14) post tamoxifen. #p<0.001 vs. controls. One-way ANOVA, Tukey’s post hoc. e, In the same animals that received cutaneous stimulation, muscle mechanical thresholds increased over time in all groups. *p<0.001 vs. BL, Two-way RM ANOVA, Tukey’s post hoc. e’, Area under the curve analysis for muscle mechanical thresholds in combined groups indicates a reduced threshold in GHr mutant groups compared to controls. #p=0.002 vs. controls. One-way ANOVA, Tukey’s post hoc. f, Dynamic muscle strength indicated that all groups except Adv;GHr^f/f^ increased over time by 7 days post tamoxifen. *p<0.001 vs. BL. n = same as (c). f’, Combined group area under the curve analysis reveals significantly reduced time in mutant GHr groups (n = 18-23/time point) compared to control groups (n = 65-70/time point). #p<0.001 vs. controls, one-way ANOVA, Tukey’s post hoc. Data shown as mean ± s.e.m.

### Local injection of GH at the time of neonatal muscle incision blocks behavioral hypersensitivity and primary afferent sensitization

Since peripheral GH signaling to neurons appeared to significantly influence nociception during early postnatal development, we wanted to test whether this pathway could also modulate injury-related hypersensitivity in the periphery. In order to assess this, we utilized a “reverse” hindpaw incision model in which surgical incision of the flexor digitorum brevis (FDB) muscles was obtained from the dorsal side of the foot through the hairy skin. The rationale for performing incision in this manner was to reduce cutaneous injury site effects when assessing muscle-specific pain-related behaviors (muscle squeezing). The surgical model produces robust pain-related hypersensitivity in neonates and also allowed us to deliver GH directly to the injury site. We first confirmed that a neonatal “reverse” muscle incision resulted in detectable pain-like phenotypes, while an incision only of the hairy skin did not (Extended Data Figure 2-1) similar to previous work (Brennan et al 1996) (Baccei 2016). We therefore assessed the levels of GH in the muscle after incision using western blot (Fig. 2a) and found that one day after surgery, muscle GH levels were significantly reduced. This corresponded with the observed spontaneous paw guarding behaviors and muscle mechanical hypersensitivity induced by incision one day later (Fig. 2b,c). To determine if local injection of GH could blunt incision-related hypersensitivity, we first performed a dose response analysis (Extended Data Figure 2-1) based on doses of GH that were insufficient to alter functional levels of systemic insulin-like growth factor 1 (IGF-1) (Farris et al 2007). We confirmed that a single injection of GH (1.5mg/kg in 10μL) into the incision site was able to restore muscle GH levels similar to naïve mice (Fig. 2a). Importantly, we then found that paw guarding scores and muscle mechanical withdrawal thresholds were normalized to baseline (BL) levels by this single injection of GH into the incision site compared to vehicle injected neonates with incision (Fig. 2b,c). By three days following the surgical incision, we no longer detected hypersensitivity in vehicle injected controls or GH treated animals (Fig. 2b,c). Interestingly, delayed GH treatment partially blocked spontaneous paw guarding if given within 8 hours (Fig. 2d) of surgery, possibly indicating a time sensitive application window. As a range of ages were used for these studies, we needed to determine if there were any age-related effects observed within our groups during this developmental time frame which shows changes in processes such as eye opening, hair production and cutaneous sensory responsiveness (Jankowski et al 2014) (Fitzgerald 2005) (Brust et al 2015). In analyses evaluating the distribution of ages within the P7-P14 time period, we observed no differences in injury induced nor GH treatment groups (Extended Data Figure 2-2). Analysis indicates an even distribution of data points for each group across ages indicating that no specific age resulted in the effects seen with our interventions or injury.

**Figure 2:**
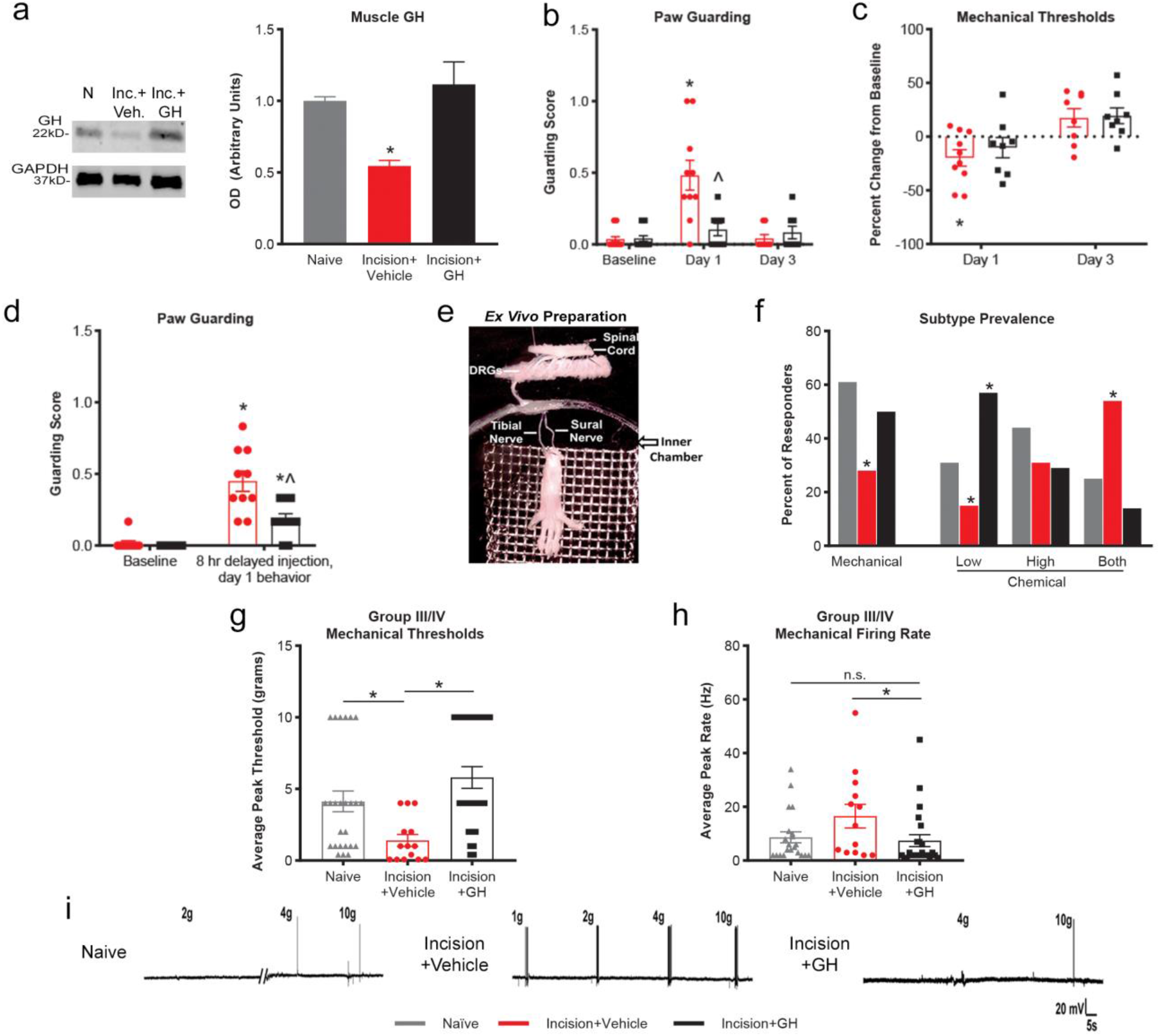
Intramuscular growth hormone injection inhibits peripheral sensitization in neonates with incision. a, Representative image and analysis of muscle growth hormone (GH) levels using western blot. Following incision, growth hormone levels are reduced in incised animals compared to naïve animals and this is restored with exogenous GH treatment. *p<0.05 vs. naïve. n = 3/group, one-way ANOVA, Tukey’s post hoc. b, Spontaneous paw guarding is increased following incision in animals treated with vehicle but not in GH treated animals at one day. By three days, neither group is different from their baseline (BL) or each other. *p<0.001 vs. BL; ^p<0.001 vs. control. n = 8-10/group, two-way RM ANOVA, Tukey’s post hoc. c, Muscle withdrawal thresholds are reduced in incised animals with vehicle injection at 1d, but this is inhibited in GH treated animals. By three days, both groups have increased withdrawal thresholds compared to baseline. *p<0.05 vs. BL. n = 8-10/group, two-way ANOVA, Tukey’s post hoc. d, When GH treatment is delayed eight hours following the injury, both groups are increased from their BL, but GH treated animals guard their paw significantly less than vehicle treated controls. *p<0.01 vs. BL, ^p<0.01 vs. control. n = 10/group, two-way RM ANOVA, Tukey’s post hoc. e, Representative image of the hind paw muscle *ex vivo* recording preparation. f, Prevalence of mechanically and chemically (“low”, “high”, or “both”) sensitive muscle afferents are altered by incision and rescued by GH treatment. *p<0.05 vs naïve, ◻^2^g, von Frey threshold of mechanically sensitive group III/IV primary afferents. Compared to naïve animals (n = 6 animals, 51 cells), surgical injury results in sensitization in incised animals (n = 8 animals, 50 cells) that is blocked by GH treatment (n = 10 animals, 54 cells). h, The average mechanical firing rate is decreased in injured animals treated with GH compared to those that were not treated after injury. *p<0.05 vs naïve. ANOVA on Ranks, Dunn’s post hoc. i, Representative mechanical responses for each group are provided. Data shown as mean ± s.e.m.

We then found that the local injection of GH at the effective dose was not sufficient to alter the cross-sectional area of individual muscle fibers or change physiological conditions of the animals including body temperature and weight. GH injection into uninjured animals also had no effects on baseline animal behaviors (Extended Data Figure 2-3). We then wanted to determine if the anti-nociceptive effects of GH treatment could be observed in older animals but found that adolescent animals (P35) do not display altered muscle growth hormone levels after an injury. Further, these animals do not show any alterations in pain-related behaviors after muscle incision in response to local GH treatment at this dose (Extended Data Figure 2-1). These data suggest that GH can modulate pain-like behaviors specifically in neonates.

Next, we used a novel neonatal *ex vivo* electrophysiological single unit recording preparation (Fig. 2e) to assess the response properties of individual primary muscle afferents in mice with incision. We found that the distribution of functional primary muscle afferents was altered by incision. Fewer mechanically sensitive group III and group IV muscle afferents (14/50, 28%) were observed in mice with muscle incision plus vehicle injection compared to naïve. A similar reduction in the numbers of innocuous metabolite (“low” responders) responsive units was also observed (2/13, 15%) that corresponded with an increase in the numbers of chemically sensitive cells that responded to both innocuous and noxious (“high” responders) metabolite mixtures (7/13, 54%). Local GH injection completely reversed the incision-related effects on muscle afferent prevalence (Fig. 2f). We also found that mechanically sensitive muscle sensory neurons had lower thresholds to mechanical deformation of the muscle receptive fields after incision plus vehicle injection, but animals treated with GH at the time of the incision did not differ in mechanical thresholds compared to naïve (Fig. 2g). GH treated animals also showed decreased firing rates (FRs) to mechanical stimuli compared to mice with incision although neither group differed from naïve (Fig. 2h,i). These parameters also were not affected by the age in the timeframe we analyzed (Extended Data Figure 2-2). The firing rates of thermally and chemically activated cells were not altered. The mean peak instantaneous frequencies were not statistically altered under any condition in any group (Extended Data Table 2-1). Taken together, exogenous local growth hormone treatment can prevent pain-related behaviors and primary afferent sensitization observed in neonates with muscle incision.

### Incision-induced transcriptional changes in the DRG can be blocked by GH treatment

To determine potential underlying neuronal mechanisms by which growth hormone mediated anti-nociception after neonatal muscle incision, we analyzed mRNA levels in the DRGs for genes previously found to be altered during neonatal injury (Jankowski et al 2014) among others known to regulate sensory responsiveness in the periphery in animals one day after injury (see full list in Table 1). Similar to that observed in mice with sensory neuron specific knockout of the GHr (see above), we found that muscle incision induced a significant increase in IGFr1 mRNA in the DRGs. Also upregulated was the proton sensor and heat transducing channel, TRPV1 and the environmental irritant and cold receptor, TRPA1. ATP sensing ion channel, P2X3 and the proton responsive channel, ASIC3, for example were not altered by incision. Interestingly, GH treatment at the time of injury blocked the upregulation of all of these factors (Table 1).

**Table 1:** Transcriptional changes following a neonatal muscle incision and local GH injection are modulated in L3/4/5 DRGs but not in the injured muscle. Data shown as a percent change from controls. n = *p<0.05 vs. controls. n = 3-12/group, one-way ANOVA with Tukey’s or ANOVA on Ranks, with Dunn’s post hoc.

| **<u>% Change in DRG mRNA</u>** |  |  |
| --- | --- | --- |
| **<u>Gene</u>** | **<u>Incision +</u>**<br>**<u>Vehicle</u>** | **<u>Incision +</u>**<br>**<u>GH</u>** |
| IGFr1 | 258 ± 29%* | 21 ± 14% |
| SRF | 142 ± 19%* | 27 ± 25% |
| TRPV1 | 178 ± 14%* | 12 ± 15% |
| TRPA1 | 493 ± 41%* | 1 ± 25% |
| STAT5 | 164 ± 20%* | 69 ± 31% |
| P2X3 | 44 ± 22%* | -2 ± 18% |
| NFκB | 83 ± 13%* | 40 ± 11% |
| ASIC3 | 18 ± 10% | 35 ± 11% |
| P2Y1 | 44 ± 15% | -14 ± 10% |
| ELK1 | -29 ± 13% | -21 ± 15% |
| ELK3 | -97 ± 5%* | -98 ± 15%* |
| ELK4 | -28 ± 8% | -19 ± 9% |
| STAT1 | 92 ± 28% | 10 ± 18% |
| STAT3 | 220 ± 7%* | 164 ± 7%* |
| IL1-r | -50 ± 15%* | -51 ± 14%* |
| TNFα-r | 15 ± 8% | 13 ± 7% |
| GHr | 15 ± 4% | 16 ± 6% |
| Fcer2a | -81 ± 20%* | -80 ± 11%* |
| OSMr | -59 ± 17%* | -66 ± 16%* |

| **<u>% Change in Muscle mRNA</u>** |  |  |
| --- | --- | --- |
| **<u>Gene</u>** | **<u>Incision +</u>**<br>**<u>Vehicle</u>** | **<u>Incision +</u>**<br>**<u>GH</u>** |
| MCP1 | 584 ± 24%* | 276 ± 17%* |
| GDNF | 204 ± 44%* | 179 ± 23%* |
| IL1β | 1098 ± 50%* | 608 ± 40%* |
| NGF | 77 ± 23%* | 72 ± 20%* |
| TNFα | 11 ± 20% | -40 ± 27% |

To assess if GH could also modulate injury related changes in gene expression in the injured muscles, we then analyzed the expression of candidate cytokines and growth factors known to be altered after injury in the periphery. Muscle incision significantly upregulated monocyte chemoattractant protein 1 (MCP-1), glial cell line-derived neurotrophic factor (GDNF), interleukin 1β (IL1β), and nerve growth factor (NGF), but not tumor necrosis factor alpha (TNFα). Interestingly, none of these factors were altered by local GH treatment (Table 1). Together, data suggests that GH provides a tonic control of gene expression in the DRG, but exogenous GH treatment at low doses may not alter the inflammatory response to incision within the muscles.

### SRF upregulation in the DRGs modulates pain-related behaviors following surgical incision

To begin to understand the mechanism by which GHr signaling effected neonatal nociception, we screened a number of known downstream transcription factors in the DRGs (Table 1). As we only see significant pain like behaviors and effects of local GH injection one day following incision, subsequent experiments evaluated this time point specifically. Neonates with muscle incision plus vehicle injection displayed significant upregulation of SRF, STAT3 and 5, and a significant downregulation of ERK like kinase 3 (ELK3), but no changes in ELK 1 or ELK4. Injection of GH into the muscles after incision specifically blocked the injury-related upregulation of SRF and STAT5. Since genetic knockout of GHr in uninjured sensory neurons regulated SRF expression (above) and to determine the role that one of these factors may play in muscle incision-induced hypersensitivity in neonates, we utilized our *in vivo* nerve-targeted siRNA knockdown strategy to inhibit the DRG upregulation of SRF (Jankowski et al 2009) (Liu et al 2017). Prior to incision, animals were injected with Penetratin-linked siRNAs against SRF (PenSRF) into the right sciatic nerve. We found that this strategy partially, but significantly blunted the upregulation of SRF in the DRGs at the mRNA and protein levels (Fig. 3a,b). We then assessed incision-related hypersensitivity in animals with the targeted knock down of SRF and found that inhibition of this transcription factor significantly reduced spontaneous paw guarding compared to neonates with control siRNA injection (PenCON) plus incision (Fig. 3c). Additionally, incision-related muscle mechanical hypersensitivity observed in mice with PenCON injection was also inhibited in mice with SRF knockdown (Fig. 3d). Interestingly, while PenSRF inhibited both guarding and mechanical hypersensitivity in males, female guarding was unaffected by PenSRF injection. Age distribution also showed no obvious effects in either group (Extended Data Figure 3-1). These behavioral changes corresponded with inhibition of TRPA1 upregulation after SRF knockdown DRGs (PenCON = 493 ± 41%*, p<0.05, PenSRF = 52 ± 37%, p>0.05; vs. naïve, n = 3-6/group). Interestingly, TRPV1 upregulation was not altered (PenCON = 90 ± 16%*, PenSRF = 128 ± 13%*, p<0.05 vs. naïve, n = 3-6/group). Together, these data indicate one neuronal transcriptional pathway downstream of GHr that may modulate neonatal nociception.

**Figure 3:**
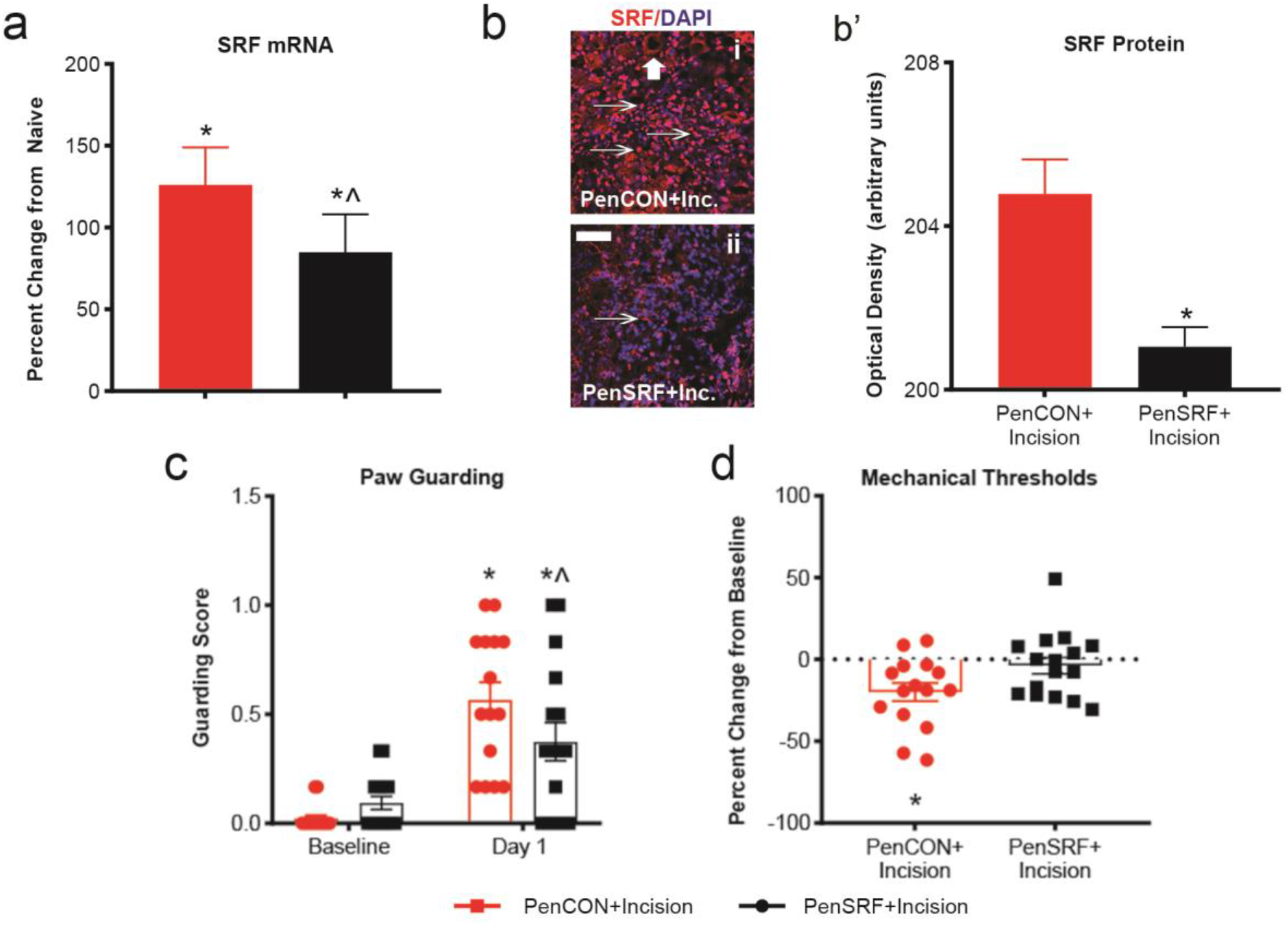
Nerve-targeted knockdown of SRF upregulation blunts pain-like responses after muscle incision. a, Injection if SRF targeting siRNAs (PenSRF) into the sciatic nerve reduces the upregulation of SRF mRNA in the DRGs after hind paw incision compared to incised mice injected with control siRNAs (PenCON). *p<0.001 vs. naïve, ^p<0.05 vs. PenCON. n = 7 naïve, 22 PenCON, 19 PenSRF, one-way ANOVA, Holm-Sidak post hoc. b, Representative images (bi-ii) display similar results at the protein level, arrows indicate SRF+ staining, big arrow indicates satellite cell, (b’) using immunocytochemical labeling and mean staining value of a ROI (red). DAPI (blue) co-stain was used to mark all nuclei. *p<0.05 vs. PenCON. n = 3 animals/group, Tukey’s post hoc. c, Incised animals show increased paw guarding over time, and animals injected with PenSRF showed reduced guarding compared to the PenCON injected mice. d, PenCON incised animals have reduced mechanical withdrawal thresholds compared to baseline, and PenSRF animals display significantly less reduction compared to PenCON injected animals. *p<0.05 vs BL, ^p<0.05 vs. PenCON. n = 15-16/group, two-way RM ANOVA, with Tukey’s post hoc. Data shown as mean ± s.e.m or percent change from naïve.

### GH sequestering by infiltrating macrophages regulates incision-related hypersensitivity in neonates

In our current and previous reports (Liu et al 2017), we have shown that the levels of GH decrease in the injured tissue within one day. However, the mechanism behind this reduction in GH levels were previously unknown. Using immunohistochemical analyses, we observed that GH is normally found in a diffuse pattern between myofibers within the muscles of naïve neonates (Fig. 4a). However, after incision, we found that GH was displaced in the muscle tissue into a more concentrated, localized manner (Fig. 4b). The areas of intense GH staining appeared to be monocyte-like based on qualitative morphological assessments. We therefore used a transgenic mouse line in which a tdTomato reporter was expressed in monocytes and mature macrophages (LysM-Cre;td-Tom) and performed immunohistochemistry for GH in these mice with incision. While non-incised hind paws have few detectable macrophages within the muscles (Extended Data Figure 4-1), animals that received a muscle incision have abundant numbers of macrophages at the injury site. Further, the concentrated pattern of GH staining in mice with incision was found to overlap significantly with the tdTomato reporter (Fig. 4c,d), while this is not observed in uninjured tissue (Extended Data Figure 4-1). Previous work has suggested that macrophages can bind and internalize GH (Govers et al 1999) (Strous et al 1996) (Lu et al 2013). Since tissue collection for our western blot analyses was obtained from animals with cardiac perfusion and samples for IHC were obtained from snap-frozen, fresh tissues (un-perfused), we posited that the “reduction” in muscle GH levels observed after incision were due to macrophage dependent sequestering. We found that following cardiac perfusion with ice cold saline in mice with incision, macrophages were no longer detected within the muscle. Moreover, in un-perfused neonates, there was no detectable reduction in GH using western blot analysis (Extended Data Figure 4-1).

**Figure 4:**
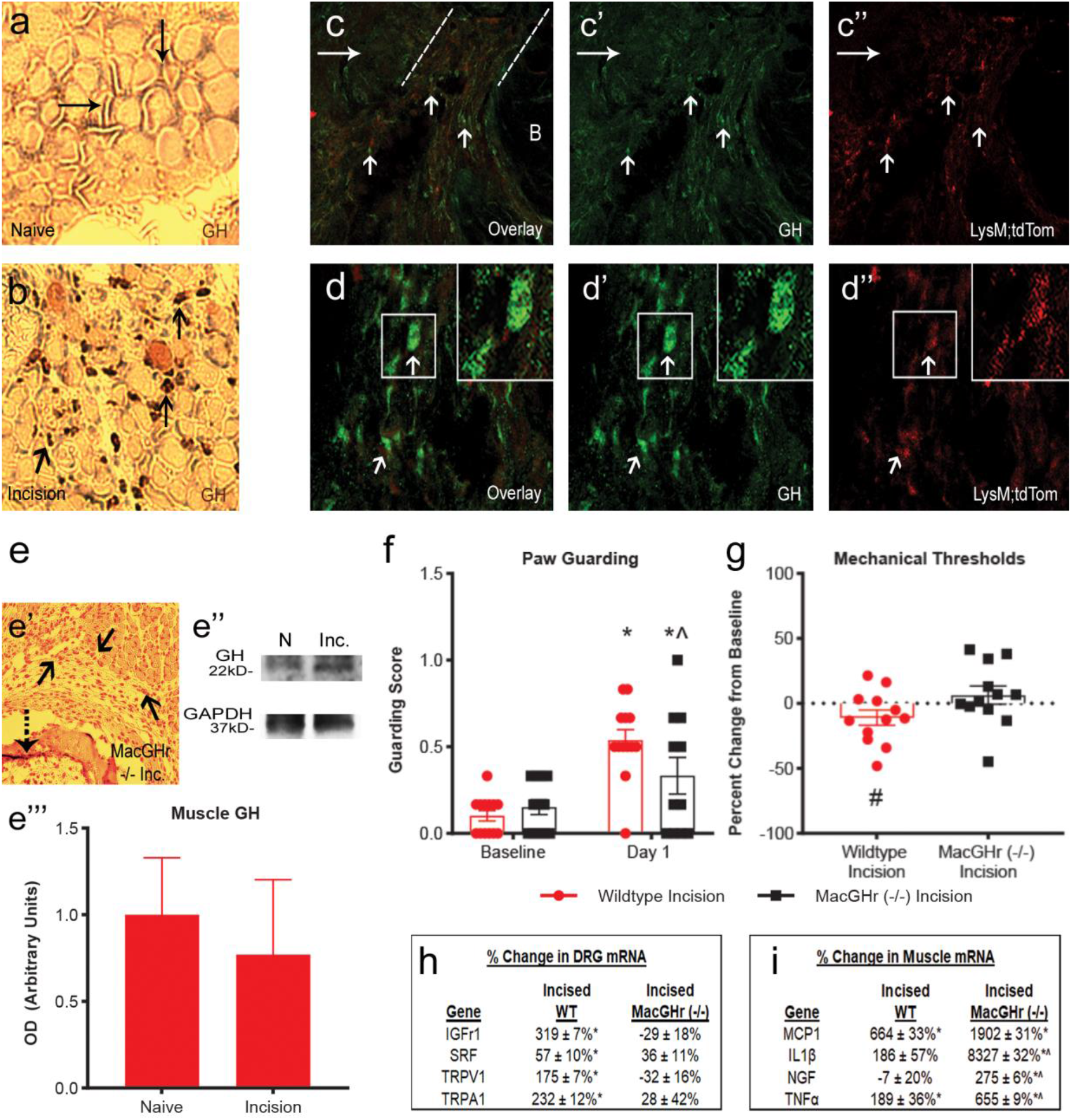
Muscle growth hormone distribution is altered by injury and macrophage specific GHr knockout attenuates pain-like phenotypes. a, Immunostaining for GH (arrows) in a cross section of hindpaw muscle of a naïve animal with DAB (3,3′-Diaminobenzidine) staining. b, GH staining (arrows) in the muscle of an animal 1d after an incision. c, Overlay of macrophages and GH (arrows) in the muscles of a LysMcre/tdTomato reporter animal using an antibody stain for GH (green) (c’) and tdTom reporter for macrophages (red) (c’’) one day after an injury of the hindpaw muscle. d-d’’ Representative higher magnification images of macrophage and GH overlay (arrows) in injured muscle. e, Representative image of GH staining (dashed arrow) using DAB in the muscles of an injured macrophage specific growth hormone receptor knockout (MacGHr−/−) animal. Hematoxylin (light red) dense cells show infiltrating macrophages (solid arrows). e’-e’’, Representative western blots and quantification of growth hormone levels after injury in MacGHr−/− animals. f, MacGHr knockout animals display less guarding compared to WT controls at 1d after injury, though both groups display increased guarding from baseline. g, MacGHr−/− mice do not show reduced muscle withdrawal thresholds after injury while WT controls display decreased thresholds. *p<0.05 vs. BL, ^p<0.05 vs. controls, #p=0.076 vs. BL. n = 12 - 13/group, one-way or two-way ANOVA, with Tukey’s post hoc. h, Upregulation of select receptors/channels in the DRGs are observed in WT control mice with incision, but this is not found in incised MacGHr−/− DRGs. i, Incision-induced upregulation of select cytokines and growth factors in the muscles is found in both control and MacGHr−/− mice. *p<0.05 vs. naïve, ^p<0.05 vs. WT incised. n = 3-4/group, two-way ANOVA, Tukey’s post hoc. Data shown as mean ± s.e.m or percent change from naïve.

Data thus suggested that under neonatal injury conditions, infiltrating macrophages may sequester GH and thereby effectively remove the tonic signal that GH normally provides to innervating primary afferents. To test this hypothesis, we first treated incised animals intramuscularly with clodronate liposomes to deplete the infiltrating macrophages. Following surgery, this depletion prevented evoked muscle hypersensitivity but not spontaneous paw guarding behaviors (Extended Data Figure 4-2). As macrophages appeared to at least play some role in incision-related hypersensitivity in neonates, we then analyzed incised animals with the GHr knocked out in macrophages and mature monocytes (LysM-Cre;GHr^f/f^; MacGHr−/−). Our injury experiments up to this point were completed in Swiss Webster male and female animals. As the MacGHr−/− mice were bred on a C57Bl/6 background, we compared the two strains in both males and females and found no difference in pain-like behaviors between strains. (Extended Data Figure 4-2). Following this, we then found that animals with the GHr knocked out in macrophages still display infiltration of these immune cells within the muscles after incision, but they do not contain GH (Fig. 4e). We also found no reduction in muscle GH levels in these mice after injury and perfusion, unlike that observed in WT animals (Fig. 4e). The MacGHr−/− mice also displayed significantly less paw guarding (Fig. 4f) and did not exhibit reduced muscle withdrawal thresholds one day after injury (Fig. 4g) unlike control mice. Mechanical hypersensitivity from the injury was not detected at 3d in either group, but unlike MacGHr−/− mice, controls still displayed a small guarding response at 3d (Extended Data Figure 4-2). Similar to other conditions, no effect of age within the P7-P14 time period was detected under these conditions (Extended Data Figure 4-3).

We also analyzed the transcriptional changes within DRGs and the injured muscles from these mice. Receptors and channels shown previously (Table 1) to be upregulated in animals after injury were also observed in control mice with incision, but these same receptors were not upregulated in MacGHr−/− animals with incision (Fig. 4h). However, the upregulation of muscle cytokines and growth factors observed in control animals with incision were unaffected by knockout of GHr in macrophages. In fact, the levels of some of these factors were significantly greater in MacGHr−/− mice with incision compared to incised controls (Fig. 4i). These data indicate that macrophages may sequester muscle GH after injury, likely removing the tonic inhibition that GH provides on primary afferent neurons and leads to injury-induced pain-related behaviors.

### Early life growth hormone reduction regulates neonatal “priming” of nociceptive responses to injury later in life

To determine if GH-related anti-nociception could modulate the prolonged effects of repeated injury, we first assessed whether GH deficiency alone could induce a prolonged hypersensitivity after incision. In order to test this, we used the GH releasing hormone receptor (GHRHr) knockout mice that we have previously shown to display neonatal specific hypersensitivity (Ford et al 2019) and performed a single incision in these mice at P35. At baseline (BL), we detected no difference between WT controls (GHRHr+/+), heterozygous GHRHr mutants (GHRHr+/−) and the homozygous GHRHr knockout (GHRHr−/−) mice. However, GHRHr−/− animals and to a degree, GHRHr+/− mice, displayed prolonged spontaneous paw guarding (Fig. 5a) and muscle withdrawal thresholds (Fig. 5b) after P35 incision compared to WT C57Bl/6 controls. In Swiss Webster animals we then confirmed previous reports (Walker et al 2009) (Moriarty et al 2018) that an early life muscle incision (P7) in WT control mice followed by a P35 incision resulted in a longer lasting mechanical hypersensitivity compared to mice with P35 incision alone. Interestingly, if treatment with GH was given at the time of the early life injury (P7), prolonged hypersensitivity was not detected after the second incision (Fig. 5c). These data support our earlier findings that GHr signaling in early life is important for sensitization and disruptions in this signaling can induce long term somatosensory alterations in response to injury later in life.

**Figure 5:**
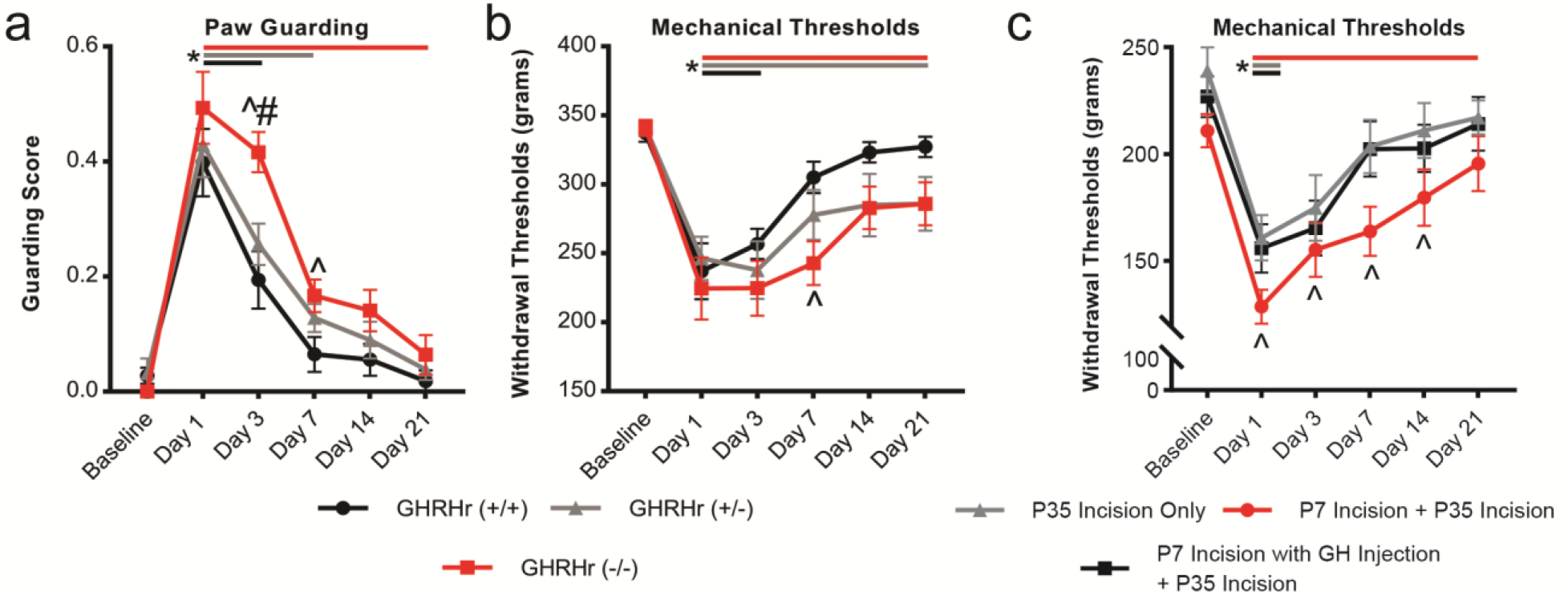
Alterations in GH levels during early postnatal development modulate the behavioral responses to incision in adolescence. a, GHRHr−/− animals guard their injured limb to a greater degree and for a longer time after P35 incision compared to GHRHr+/− and WT controls (GHRHr+/+). b, Muscle mechanical hypersensitivity is also more severe and prolonged in GHRHr+/− and GHRHr−/− mice with incision at P35 compared to GHRHr+/+ (WT C57/Bl6) animals. *p<0.05 vs. BL (indicated for each group by the colored horizontal bars next to the “*” for each group, the ending of the bar indicates the ending of the detected significance); ^p<0.05 vs. time-matched GHRHr+/+; #p<0.001 vs. time-matched GHRHr+/−. n = 9-13/group, two-way RM ANOVA with Tukey’s or ANOVA on Ranks with Dunn’s post hoc. c, Muscle withdrawal thresholds in wildtype Swiss Webster mice following a P7 plus P35 incision are prolonged compared to mice that only received a single injury at P35. This effect is blocked by injection of GH during P7 incision. *p<0.05 vs. BL (indicated as in “a” and “b”. ^p<0.05 vs. time-matched P35 incision only. n = 11-12/group, two-way RM ANOVA, with Tukey’s post hoc. Data shown as the mean ± s.e.m.

## Discussion

Our data indicate that in neonates, growth hormone expressed throughout the hindpaw muscles tonically activates the growth hormone receptor on neurons. This maintains basal transcription factor levels in sensory neurons, which supports normal afferent function. However, after an early life injury, macrophage infiltration results in an immune cell-dependent sequestration of muscle GH. The displacement of GH within the muscle removes GH signaling to neurons resulting, at least in part, in SRF-dependent transcription of sensory-related receptors/channels in the affected DRGs. Restoring the basal levels of GH (through targeted injections or by preventing the macrophage dependent sequestration), maintains neuronal GHr signaling and blocks peripheral sensitization (Fig. 6). GH signaling to immature neurons also appears to modulate injury responses later in life (Fig. 5). Thus, in neonates, a unique immune system dependent modulation of endocrine communication with the peripheral nervous system is accessed after injury to regulate nociception.

**Figure 6:**
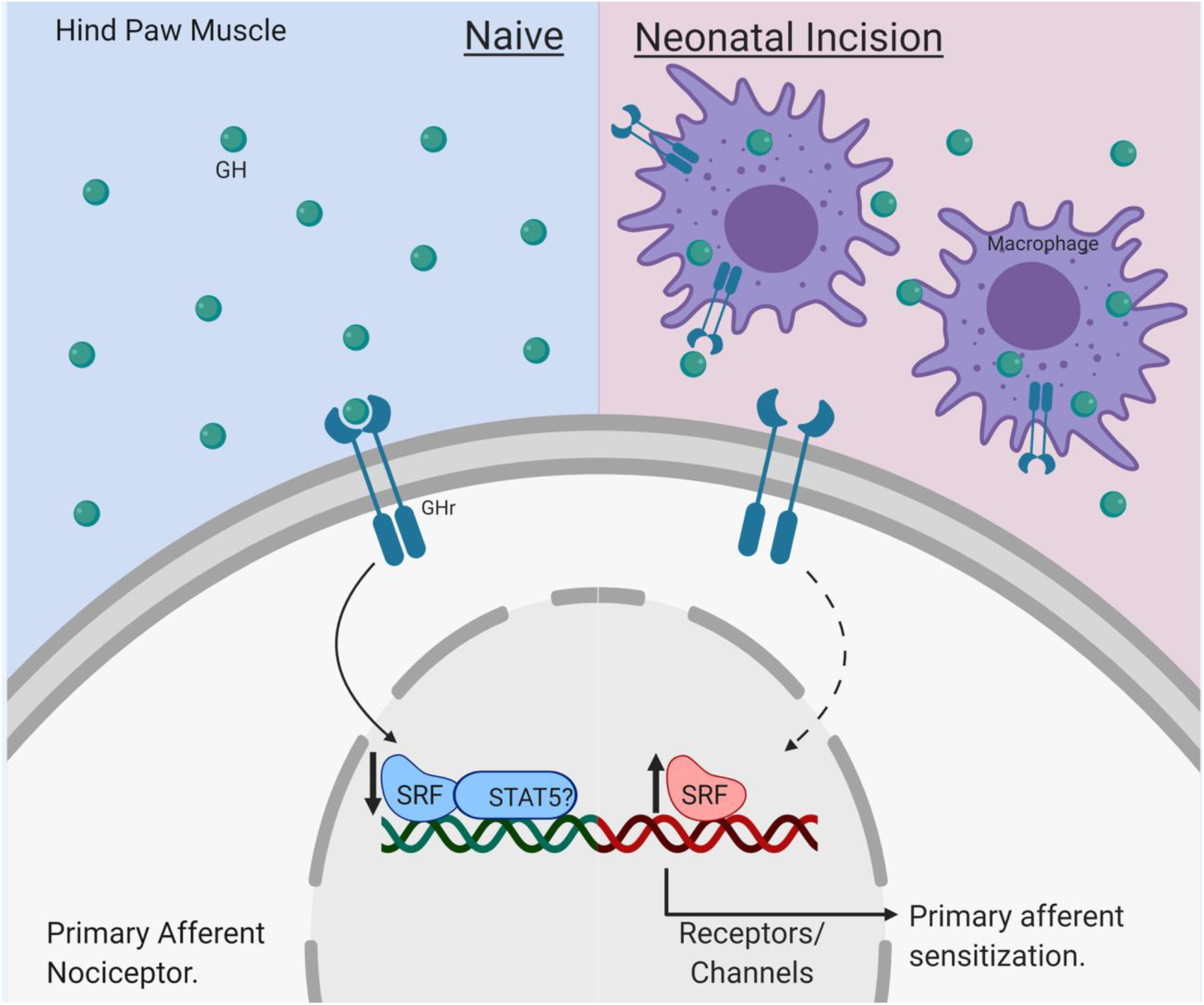
Role of GH in neonatal nociception. Under naïve conditions (left), GH is diffusely available throughout the neonatal muscle tissue, allowing persistent activation of the GHr on primary afferent nociceptors. Normal activation of neuronal GHr maintains homeostatic transcription levels and regulates sensitizing factors such as TRPA1 and TRPV1 partially through SRF-dependent regulation. Following a neonatal injury (right), macrophages infiltrate the injured tissue and sequester GH, thereby effectively reducing GH-availability to the nociceptors. The removal of tonic GH signaling to neurons permits SRF (and possibly STAT5) dependent transcription of various receptors and channels, leading to nociceptor sensitization and pain-related hypersensitivity after injury.

Nociception involves a variety of distinct extracellular and intracellular neuronal mechanisms, each of which can play a unique role in how peripheral stimuli are perceived upon transmission to the CNS (Julius & Basbaum 2001). Throughout development, distinct neurotrophic factors regulate neuronal subtype formation and innervation; however, these growth factors also display non-canonical roles (Moqrich et al 2004) (Shelton & Reichardt 1984) (Mendell 1996). Recent investigations also demonstrate that developing sensory neurons display changing transcriptional identities and functional maturation from the neonatal period through adulthood (Sharma et al 2020) (Jankowski et al 2014) (Adelman et al 2019). While growth factors clearly alter subtype survival, we detect that modulation of neuronal GH signaling results in functional changes to normal somatosensory processing in both the skin (Ford et al 2019) and the muscle. GH may have a more global effect on afferent function that influences somatosensory development unique to that observed with neurotrophic factors.

Recent data also suggests that the developmental stage of the animal may affect how primary afferents respond to injury (Jankowski et al 2014). Our previous and current reports however indicate that muscle afferent sensitization occurs similarly across the lifespan but may become insensitive to GH as an antinociceptive treatment beginning in adolescence (Extended Data Figures 2-1, 2-2, 3-2, 4-3) (Queme et al 2020). Further, the developing immune system shows unique characteristics resulting in a microenvironment that is distinct from that of older subjects (Kumar & Bhat 2016) (Winterberg et al 2015) (Weston et al 1977). Here, we show that GH may have specific roles in regulating multiple cell types in the periphery after insult. Due to the unique developmental functions of primary afferents and the immune systems, neonatal injury may engage a distinct cellular interaction through which GH ultimately modulates neonatal nociception and injury-related hypersensitivity (Figs. 2–6). It is worth noting that the ages we detect GH-related effects on nociception are also when systemic GH is at its highest, and is critical for overall growth and development, which may account for its age-specific effects (Muller et al 1999) (Figs. 1,2; Extended Data Figure 2-1, 2-2). Together, this evidence indicates a critical period in which the nervous, immune and endocrine systems uniquely interact to modulate peripheral sensitization.

Following a noxious stimulus, primary afferents respond by initiating various intracellular cascades that result in an upregulation of factors that change how the neuron responds to specific sensory stimuli (Basbaum et al 2009). In our current work, we found that neonatal incision upregulated SRF and STAT5 in the DRGs, which could be blocked by intramuscular GH treatment (Fig. 2; Table 1). However, only SRF was found to be altered in the DRGs in our sensory neuron GHr knockout animal (Fig. 1). SRF is a well-studied transcription factor that operates in a complex within the nucleus to regulate gene expression in an age-controlled manner (Zhang et al 2014). Further, SRF is a downstream target of canonical GH signaling (Carter-Su et al 2016). Although we did not directly target this factor in the Adv;GHr^f/f^ mice, we were able to show that incision-induced upregulation of SRF is important for at least TRPA1, but not TRPV1 upregulation after incision. STAT5 is another interesting target that is regulated by muscle GH signaling to neurons after injury, but not GHr signaling directly. It is possible that STAT5 has the ability to transcriptionally control TRPV1 (TargetScanMouse and miRDB), but that upregulation of this transcription factor under neonatal injury conditions is due to non-GHr related signaling pathways (Gouin et al 2017) (Kao et al 2012). It is likely that a number of alterations are induced by both injury and GH signaling resulting in an intracellular dynamic that collectively modulates sensitization. Further analyses will be necessary to determine this.

Immune system regulation of the affected microenvironment is a well-known phenomenon following many different injuries (Basbaum et al 2009) (Julius & Basbaum 2001). The peripheral immune system reacts to an injury by mobilizing in stages with macrophages being one of the first responders (Fig. 4). This effect may be more heavily relied upon in neonates that have yet to fully develop (Winterberg et al 2015). We detected a time-dependent window for the local injection of GH that, if given 8 hours after the injury, only partially blocks sensitization (Fig. 2d). It may be that this time point corresponds with the infiltration of macrophages to the injured area and is belatedly able to restore muscle GH levels for nociceptor signaling prior to full sensitization. Further, previous reports indicate that the use of free GH by macrophages is necessary for proper immune function such as control over their production of cytokines and growth factors which subsequently regulates the inflammatory response to injury (Lu et al 2013). Here, we found that the prevention of GH signaling to macrophages by specific deletion of the GHr in these cells resulted in a reduced control over injury-site specific cytokine levels (Fig. 4) (Lu et al 2013) (Kumar et al 2014). It is worth noting that recent work has indicated an opposite effect, demonstrating that activating macrophages in culture with GH also results in an increased pro-inflammatory cytokine release (Schneider et al 2019). It is likely that GH signaling is used by these effector cells as a homeostatic regulator (Dehkhoda et al 2018). Despite the increase of these factors known to be pro-nociceptive (Sommer & Kress 2004), our manipulation, which resulted in rescued muscle GH levels, prevented sensitization and pain-like behaviors. Interestingly, we also detected that the receptors of some cytokines, such as the IL1β receptor, are downregulated in DRG after a neonatal injury which may account for this discrepancy (Table 1). Also, it may be that restoration of GH signaling in neurons supersedes pro-nociceptive cytokine signaling (Figs. 1, 2, 5). It will be important to test this notion in the future.

Early life injury induces neuronal and immunological alterations that, when re-activated by an insult later in life, results in enhanced pain-like outcomes (Ren et al 2004) (Boissé et al 2005) (Walker et al 2009) (Schwaller et al 2015) (Moriarty et al 2019) (Zhong et al 2018). While a number of analyses of the central nervous and immune systems have confirmed that this “priming” effect requires central processing, it is clear that primary afferent input is also necessary (Moriarty et al 2019) (Beggs et al 2012) (Walker et al 2016) (Moriarty et al 2018). As our current (Fig. 2) and previous work (Liu et al 2017) suggested that peripheral GH signaling regulated afferent sensitization and pain-related behaviors after injury, we posited that GH may also influence the “priming” effects that neonatal injury has on adolescent responses to re-injury. GH deficiency which induces a neonatal specific hypersensitivity to peripheral stimuli (Ford et al 2019) is sufficient to prolong the normal behavioral responses to surgical insult similar to the repeated injury model (Fig. 5). This data is supported by recent findings that shows in a different model of growth hormone deficiency, that animals are overall similar at adult baseline levels, however, are hypersensitive following a hindpaw formalin challenge (Leone et al 2019). These data suggest that GH levels and immune modulation are important for the induction of the early life injury “priming” effect and that this system can be manipulated by exogenous GH treatment.

Here, we describe a potentially neonatal specific mechanism of nociception as well as an intervention that is effective at blocking pain-like behaviors and primary afferent sensitization. Correlations between the level of circulating GH levels and pain in patients has been previously observed in patients deficient in the hormone (Cimaz et al 2001). Interestingly, patients that suffer from widespread muscle pain (fibromyalgia) sometimes also have altered GH levels and can be treated with GH for pain. These results may have important clinical implications as many children are not diagnosed with growth hormone deficiency until they are a few years of age, in which time they may have already induced a “priming” effect to later in life injuries such as surgery. Together with the current and previous work (Liu et al 2017), data suggest that interventions designed to control the local levels of GH may be clinically beneficial for pain in young children.

## Author contributions

A.J.D. and M.P.J. conceived and designed the study and completed the *ex vivo* experiments and analysis. A.J.D. collected most of the data and completed the analyses. Z.K.F., K.J.G. and C.E.M. collected 1d MacGHrKO, clodronate, some GH-treatment, GHBP, and WT dual incision data. M.H. and R.C.H. managed the colony and performed genotyping assays in addition to performing some immunohistochemical and cell culture experiments.

## Acknowledgements

This work was supported by grants to MPJ from the NIH (R01NS105715, R56NS103178) and the Rita Allen Foundation in addition to support from the Cincinnati Children’s Research Foundation and Department of Anesthesia. We would also like to thank Dr. Ram Menon from the University of Michigan for supplying us with the macrophage specific GH receptor knockout mice.

## References

Adelman P, Baumbauer K, Friedman R, Shah M, Wright M, et al. 2019. Single cell q-PCR derived expression profiles of identified sensory neurons. bioRxiv: 560672

Au - Feather-Schussler DN, Au - Ferguson TS. 2016. A Battery of Motor Tests in a Neonatal Mouse Model of Cerebral Palsy. JoVE: e53569

Baccei ML. 2016. Rewiring of Developing Spinal Nociceptive Circuits by Neonatal Injury and Its Implications for Pediatric Chronic Pain. Children 3: 16

Baoge L, Van Den Steen E, Rimbaut S, Philips N, Witvrouw E, et al. 2012. Treatment of skeletal muscle injury: a review. ISRN orthopedics 2012: 689012

Bartholomew EF MF, Nath JL. 2009. Fundamentals of anatomy & physiology. Upper Saddle River, NJ: Pearson Education Inc. 616–617

Basbaum AI, Bautista DM, Scherrer G, Julius D. 2009. Cellular and molecular mechanisms of pain. Cell 139: 267–84

Beggs S, Currie G, Salter MW, Fitzgerald M, Walker SM. 2012. Priming of adult pain responses by neonatal pain experience: maintenance by central neuroimmune activity. Brain 135: 404–17

Boissé L, Spencer SJ, Mouihate A, Vergnolle N, Pittman QJ. 2005. Neonatal immune challenge alters nociception in the adult rat. Pain 119: 133–41

Brennan TJ, Vandermeulen EP, Gebhart GF. 1996. Characterization of a rat model of incisional pain. Pain 64: 493–501

Brewer CL, Baccei ML. 2019. The development of pain circuits and unique effects of neonatal injury. Journal of neural transmission (Vienna, Austria : 1996)

Brust V, Schindler PM, Lewejohann L. 2015. Lifetime development of behavioural phenotype in the house mouse (Mus musculus). Frontiers in Zoology 12: S17

Carter-Su C, Schwartz J, Argetsinger LS. 2016. Growth hormone signaling pathways. Growth Horm IGF Res 28: 11–5

Ceseña TI, Cui TX, Piwien-Pilipuk G, Kaplani J, Calinescu AA, et al. 2007. Multiple mechanisms of growth hormone-regulated gene transcription. Molecular genetics and metabolism 90: 126–33

Cimaz R, Rusconi R, Fossali E, Careddu P. 2001. Unexpected healing of cutaneous ulcers in a short child. Lancet (London, England) 358: 211–2

Cuatrecasas G. 2009. Fibromyalgic syndromes: could growth hormone therapy be beneficial? Pediatric endocrinology reviews : PER 6 Suppl 4: 529–33

Cuatrecasas G, Alegre C, Casanueva FF. 2014. GH/IGF1 axis disturbances in the fibromyalgia syndrome: is there a rationale for GH treatment? Pituitary 17: 277–83

Cuatrecasas G, Alegre C, Fernandez-Solà J, Gonzalez MJ, Garcia-Fructuoso F, et al. 2012. Growth hormone treatment for sustained pain reduction and improvement in quality of life in severe fibromyalgia. Pain 153: 1382–9

Cuatrecasas G, Gonzalez MJ, Alegre C, Sesmilo G, Fernandez-Solà J, et al. 2010. High prevalence of growth hormone deficiency in severe fibromyalgia syndromes. The Journal of clinical endocrinology and metabolism 95: 4331–7

Cuatrecasas G, Riudavets C, Güell MA, Nadal A. 2007. Growth hormone as concomitant treatment in severe fibromyalgia associated with low IGF-1 serum levels. A pilot study. BMC musculoskeletal disorders 8: 119

Dallman MA, Ladle DR. 2013. Quantitative analysis of locomotor defects in neonatal mice lacking proprioceptive feedback. Physiology & behavior 120: 97–105

Dattani M, Preece M. 2004. Growth hormone deficiency and related disorders: insights into causation, diagnosis, and treatment. Lancet (London, England) 363: 1977–87

Dehkhoda F, Lee CMM, Medina J, Brooks AJ. 2018. The Growth Hormone Receptor: Mechanism of Receptor Activation, Cell Signaling, and Physiological Aspects. Front Endocrinol (Lausanne) 9: 35

Dubick MN, Ravin TH, Michel Y, Morrisette DC. 2015. Use of localized human growth hormone and testosterone injections in addition to manual therapy and exercise for lower back pain: a case series with 12-month follow-up. Journal of Pain Research 8: 295–302

Farris GM, Miller GK, Wollenberg GK, Molon-Noblot S, Chan C, Prahalada S. 2007. Recombinant rat and mouse growth hormones: risk assessment of carcinogenic potential in 2-year bioassays in rats and mice. Toxicological sciences : an official journal of the Society of Toxicology 97: 548–61

Fitzgerald M. 2005. The development of nociceptive circuits. Nature Reviews Neuroscience 6: 507–20

Ford ZK, Dourson AJ, Liu X, Lu P, Green KJ, et al. 2019. Systemic growth hormone deficiency causes mechanical and thermal hypersensitivity during early postnatal development. IBRO Reports 6: 111–21

Gomez R, Kohler DM, Brackley AD, Henry MA, Jeske NA. 2018. Serum response factor mediates nociceptor inflammatory pain plasticity. Pain reports 3: e658

Goodrich CA. 1977. Measurement of body temperature in neonatal mice. Journal of applied physiology: respiratory, environmental and exercise physiology 43: 1102–5

Gouin O, L’Herondelle K, Lebonvallet N, Le Gall-Ianotto C, Sakka M, et al. 2017. TRPV1 and TRPA1 in cutaneous neurogenic and chronic inflammation: pro-inflammatory response induced by their activation and their sensitization. Protein & cell 8: 644–61

Govers R, ten Broeke T, van Kerkhof P, Schwartz AL, Strous GJ. 1999. Identification of a novel ubiquitin conjugation motif, required for ligand-induced internalization of the growth hormone receptor. The EMBO journal 18: 28–36

Hermann C, Hohmeister J, Demirakca S, Zohsel K, Flor H. 2006. Long-term alteration of pain sensitivity in school-aged children with early pain experiences. Pain 125: 278–85

Hester MS, Danzer SC. 2013. Accumulation of abnormal adult-generated hippocampal granule cells predicts seizure frequency and severity. J Neurosci 33: 8926–36

Jankowski MP, McIlwrath SL, Jing X, Cornuet PK, Salerno KM, et al. 2009. Sox11 transcription factor modulates peripheral nerve regeneration in adult mice. Brain research 1256: 43–54

Jankowski MP, Ross JL, Weber JD, Lee FB, Shank AT, Hudgins RC. 2014. Age-dependent sensitization of cutaneous nociceptors during developmental inflammation. Mol Pain 10: 34

Julius D, Basbaum AI. 2001. Molecular mechanisms of nociception. Nature 413: 203–10

Kao DJ, Li AH, Chen JC, Luo RS, Chen YL, et al. 2012. CC chemokine ligand 2 upregulates the current density and expression of TRPV1 channels and Nav1.8 sodium channels in dorsal root ganglion neurons. Journal of neuroinflammation 9: 189

Koerber HR, McIlwrath SL, Lawson JJ, Malin SA, Anderson CE, et al. 2010. Cutaneous C-polymodal fibers lacking TRPV1 are sensitized to heat following inflammation, but fail to drive heat hyperalgesia in the absence of TPV1 containing C-heat fibers. Mol Pain 6: 58

Koerber HR, Woodbury CJ. 2002. Comprehensive phenotyping of sensory neurons using an ex vivo somatosensory system. Physiology & behavior 77: 589–94

Kumar PA, Chitra PS, Lu C, Sobhanaditya J, Menon R. 2014. Growth hormone (GH) differentially regulates NF-kB activity in preadipocytes and macrophages: implications for GH’s role in adipose tissue homeostasis in obesity. Journal of physiology and biochemistry 70: 433–40

Kumar SK, Bhat BV. 2016. Distinct mechanisms of the newborn innate immunity. Immunology letters 173: 42–54

Lau J, Minett MS, Zhao J, Dennehy U, Wang F, et al. 2011. Temporal control of gene deletion in sensory ganglia using a tamoxifen-inducible Advillin-Cre-ERT2 recombinase mouse. Mol Pain 7: 100

Leone S, Chiavaroli A, Recinella L, Orlando G, Ferrante C, et al. 2019. Increased pain and inflammatory sensitivity in growth hormone-releasing hormone (GHRH) knockout mice. Prostaglandins & other lipid mediators: 106362

Lim Y, Godambe S. 2017. Prevention and management of procedural pain in the neonate: an update, American Academy of Pediatrics, 2016. Archives of disease in childhood. Education and practice edition 102: 254–56

Liu X, Green KJ, Ford ZK, Queme LF, Lu P, et al. 2017. Growth hormone regulates the sensitization of developing peripheral nociceptors during cutaneous inflammation. Pain 158: 333–46

Lu C, Kumar PA, Sun J, Aggarwal A, Fan Y, et al. 2013. Targeted deletion of growth hormone (GH) receptor in macrophage reveals novel osteopontin-mediated effects of GH on glucose homeostasis and insulin sensitivity in diet-induced obesity. J Biol Chem 288: 15725–35

Marsh D, Dickenson A, Hatch D, Fitzgerald M. 1999. Epidural opioid analgesia in infant rats II: responses to carrageenan and capsaicin. Pain 82: 33–8

McMahon SB, Cafferty WB, Marchand F. 2005. Immune and glial cell factors as pain mediators and modulators. Exp Neurol 192: 444–62

McMahon SB, La Russa F, Bennett DL. 2015. Crosstalk between the nociceptive and immune systems in host defence and disease. Nature reviews. Neuroscience 16: 389–402

Mendell LM. 1996. Neurotrophins and sensory neurons: role in development, maintenance and injury. A thematic summary. Philosophical transactions of the Royal Society of London. Series B, Biological sciences 351: 463–7

Moqrich A, Earley TJ, Watson J, Andahazy M, Backus C, et al. 2004. Expressing TrkC from the TrkA locus causes a subset of dorsal root ganglia neurons to switch fate. Nat Neurosci 7: 812–8

Moriarty O, Harrington L, Beggs S, Walker SM. 2018. Opioid analgesia and the somatosensory memory of neonatal surgical injury in the adult rat. British journal of anaesthesia 121: 314–24

Moriarty O, Tu Y, Sengar AS, Salter MW, Beggs S, Walker SM. 2019. Priming of Adult Incision Response by Early-Life Injury: Neonatal Microglial Inhibition Has Persistent But Sexually Dimorphic Effects in Adult Rats. The Journal of Neuroscience 39: 3081

Muller EE, Locatelli V, Cocchi D. 1999. Neuroendocrine control of growth hormone secretion. Physiological reviews 79: 511–607

Nikolaou S, Hu L, Cornwall R. 2015. Afferent Innervation, Muscle Spindles, and Contractures Following Neonatal Brachial Plexus Injury in a Mouse Model. The Journal of hand surgery 40: 2007–16

Philippou A, Maridaki M, Theos A, Koutsilieris M. 2012. Cytokines in muscle damage. Advances in clinical chemistry 58: 49–87

Pinho-Ribeiro FA, Baddal B, Haarsma R, O’Seaghdha M, Yang NJ, et al. 2018. Blocking Neuronal Signaling to Immune Cells Treats Streptococcal Invasive Infection. Cell 173: 1083–97.e22

Queme LF, Ross JL, Lu P, Hudgins RC, Jankowski MP. 2016. Dual Modulation of Nociception and Cardiovascular Reflexes during Peripheral Ischemia through P2Y1 Receptor-Dependent Sensitization of Muscle Afferents. J Neurosci 36: 19–30

Queme LF, Weyler AA, Cohen ER, Hudgins RC, Jankowski MP. 2020. A dual role for peripheral GDNF signaling in nociception and cardiovascular reflexes in the mouse. Proc Natl Acad Sci U S A 117: 698–707

Ren K, Anseloni V, Zou SP, Wade EB, Novikova SI, et al. 2004. Characterization of basal and re-inflammation-associated long-term alteration in pain responsivity following short-lasting neonatal local inflammatory insult. Pain 110: 588–96

Ren K, Dubner R. 2010. Interactions between the immune and nervous systems in pain. Nature medicine 16: 1267–76

Ross JL, Queme LF, Cohen ER, Green KJ, Lu P, et al. 2016. Muscle IL1beta Drives Ischemic Myalgia via ASIC3-Mediated Sensory Neuron Sensitization. J Neurosci 36: 6857–71

Ross JL, Queme LF, Lamb JE, Green KJ, Jankowski MP. 2018. Sex differences in primary muscle afferent sensitization following ischemia and reperfusion injury. Biology of sex differences 9: 2

Salaffi F, Giacobazzi G, Di Carlo M. 2018. Chronic Pain in Inflammatory Arthritis: Mechanisms, Metrology, and Emerging Targets-A Focus on the JAK-STAT Pathway. Pain research & management 2018: 8564215

Sass FA, Fuchs M, Pumberger M, Geissler S, Duda GN, et al. 2018. Immunology Guides Skeletal Muscle Regeneration. International journal of molecular sciences 19

Schindelin J, Arganda-Carreras I, Frise E, Kaynig V, Longair M, et al. 2012. Fiji: an open-source platform for biological-image analysis. Nat Methods 9: 676–82

Schneider A, Wood HN, Geden S, Greene CJ, Yates RM, et al. 2019. Growth hormone-mediated reprogramming of macrophage transcriptome and effector functions. Scientific reports 9: 19348

Schwaller F, Beggs S, Walker SM. 2015. Targeting p38 Mitogen-activated Protein Kinase to Reduce the Impact of Neonatal Microglial Priming on Incision-induced Hyperalgesia in the Adult Rat. Anesthesiology 122: 1377–90

Sharma N, Flaherty K, Lezgiyeva K, Wagner DE, Klein AM, Ginty DD. 2020. The emergence of transcriptional identity in somatosensory neurons. Nature 577: 392–98

Shelton DL, Reichardt LF. 1984. Expression of the beta-nerve growth factor gene correlates with the density of sympathetic innervation in effector organs. Proc Natl Acad Sci U S A 81: 7951–5

Sommer C, Kress M. 2004. Recent findings on how proinflammatory cytokines cause pain: peripheral mechanisms in inflammatory and neuropathic hyperalgesia. Neuroscience letters 361: 184–7

Strous GJ, van Kerkhof P, Govers R, Ciechanover A, Schwartz AL. 1996. The ubiquitin conjugation system is required for ligand-induced endocytosis and degradation of the growth hormone receptor. The EMBO journal 15: 3806–12

van der Lely AJ, Hutson RK, Trainer PJ, Besser GM, Barkan AL, et al. 2001. Long-term treatment of acromegaly with pegvisomant, a growth hormone receptor antagonist. Lancet (London, England) 358: 1754–9

Walker SM. 2019. Long-term effects of neonatal pain. Seminars in fetal & neonatal medicine

Walker SM, apos, Reilly H, Beckmann J, Marlow N. 2018. Conditioned pain modulation identifies altered sensitivity in extremely preterm young adult males and female. BJA: British Journal of Anaesthesia 121: 636–46

Walker SM, Beggs S, Baccei ML. 2016. Persistent changes in peripheral and spinal nociceptive processing after early tissue injury. Experimental Neurology 275: 253–60

Walker SM, Tochiki KK, Fitzgerald M. 2009. Hindpaw incision in early life increases the hyperalgesic response to repeat surgical injury: critical period and dependence on initial afferent activity. Pain 147: 99–106

Weston WL, Carson BS, Barkin RM, Slater GD, Dustin RD, Hecht SK. 1977. Monocyte-macrophage function in the newborn. American journal of diseases of children (1960) 131: 1241–2

Winterberg T, Vieten G, Meier T, Yu Y, Busse M, et al. 2015. Distinct phenotypic features of neonatal murine macrophages. European journal of immunology 45: 214–24

Ye Y, Woodbury CJ. 2010. Early postnatal loss of heat sensitivity among cutaneous myelinated nociceptors in Swiss-Webster mice. Journal of neurophysiology 103: 1385–96

Zhang X, Azhar G, Rogers SC, Foster SR, Luo S, Wei JY. 2014. Overexpression of p49/STRAP alters cellular cytoskeletal structure and gross anatomy in mice. BMC Cell Biology 15: 32–32

Zhong XS, Winston JH, Luo X, Kline KT, Nayeem SZ, et al. 2018. Neonatal Colonic Inflammation Epigenetically Aggravates Epithelial Inflammatory Responses to Injury in Adult Life. Cellular and molecular gastroenterology and hepatology 6: 65–78

