## Supplementary Information for "Early life nociception is influenced by peripheral growth hormone signaling"

**Extended Data:**

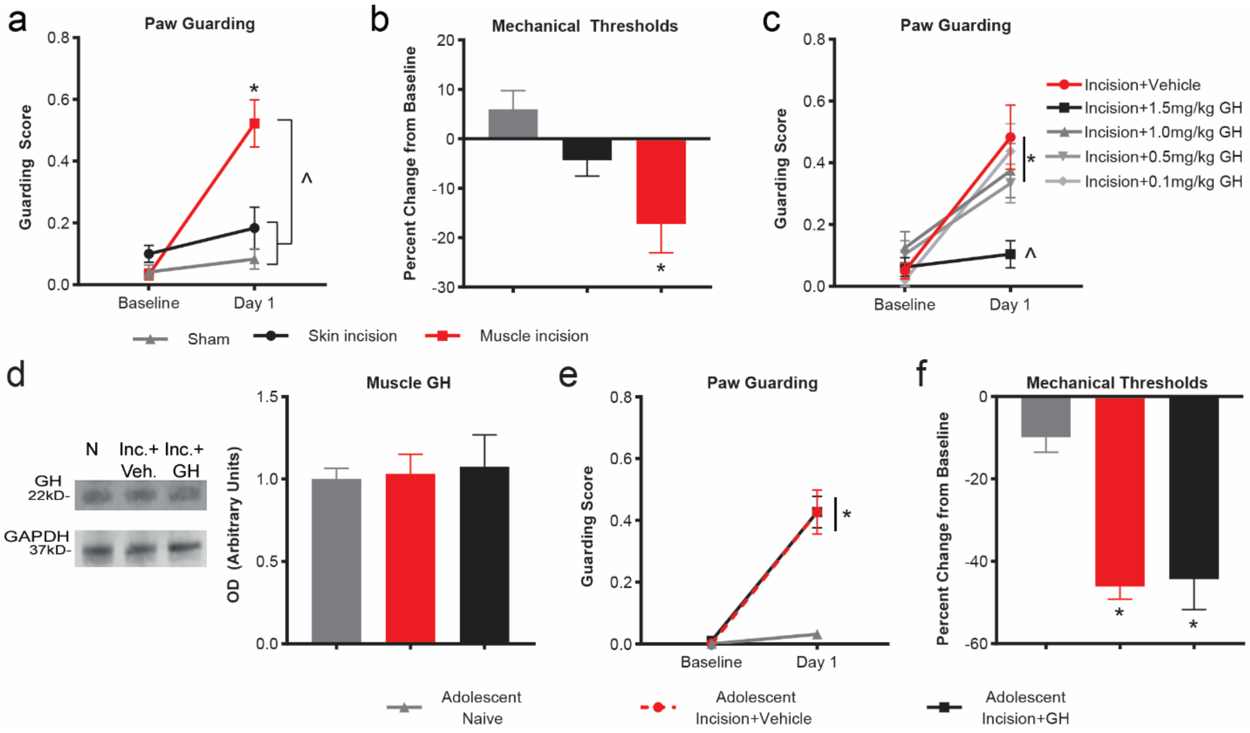


**Extended Figure 2-1: Muscle incision produces measurable behavioral responses compared to controls.** a, One day after a "reverse" neonatal muscle incision, animals guard the injured hindpaw compared to baseline and controls. Sham injured animals that received a suture through intact skin and dorsal skin incision only (not muscle) did not guard compared to baseline (BL). b, Only animals that received a muscle incision displayed lower squeezing withdrawal thresholds compared to BL. *p<0.001 vs. BL; ^p<0.001 vs. each control group. n = 10-15/group, two-way RM ANOVA, Tukey's post hoc. c, One day after an injury, animals dosed with GH at 0, 0.1, 0.5 and 1.0 mg/kg were significantly different from BL. Animals injected with 1.5 mg/kg did not differ from BL and had significantly lower guarding scores compared to vehicle injected animals at 1d. *p<0.05 vs. BL; ^p<0.001 vs. incision+vehicle. d, Representative images of western blots for GH in naïve, incised and GH treated incised animals. Quantification indicates no differences between groups. n = 4/group, one-way ANOVA. e, Adolescent animals with vehicle+incision or GH+incision guard one day after an injury compared to BL and naïve animals. f, After a hindpaw incision, both incised groups of P35 animals (vehicle and GH treated) display reduced mechanical withdrawal thresholds compared to BL and naïve. *p<0.001 vs. BL and naïve. n = 8/group, two-way RM ANOVA, Tukey's post hoc. Data shown as the mean ± s.e.m.

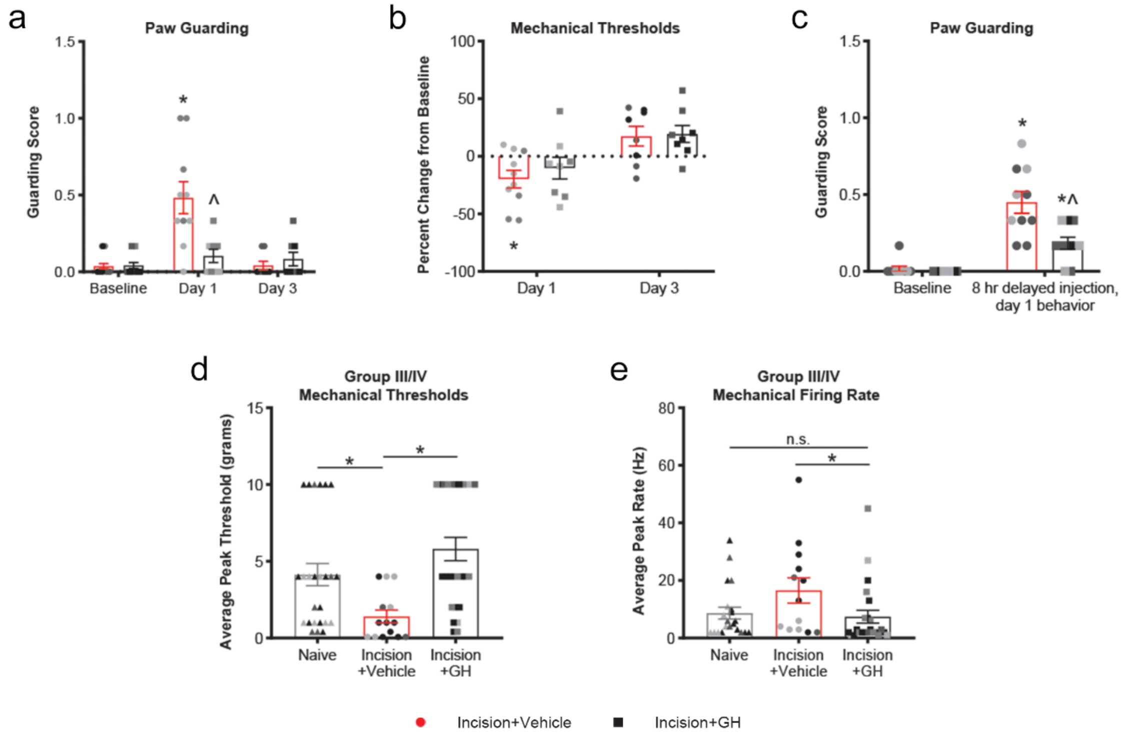


**Extended Data Figure 2-2: Age distribution for behavioral and electrophysiological data in Figure 2.** Statistics and data points are equivalent to those displayed in Figure 2 where red or gray bars indicates control group (circles or triangles) and black bars indicate experimental group (squares). Here, the shades of the data points indicate the age range of the animals. Light gray = ≤P7. Medium gray = P8 and P9. Dark gray = P10, P11. Black = ≥P12.

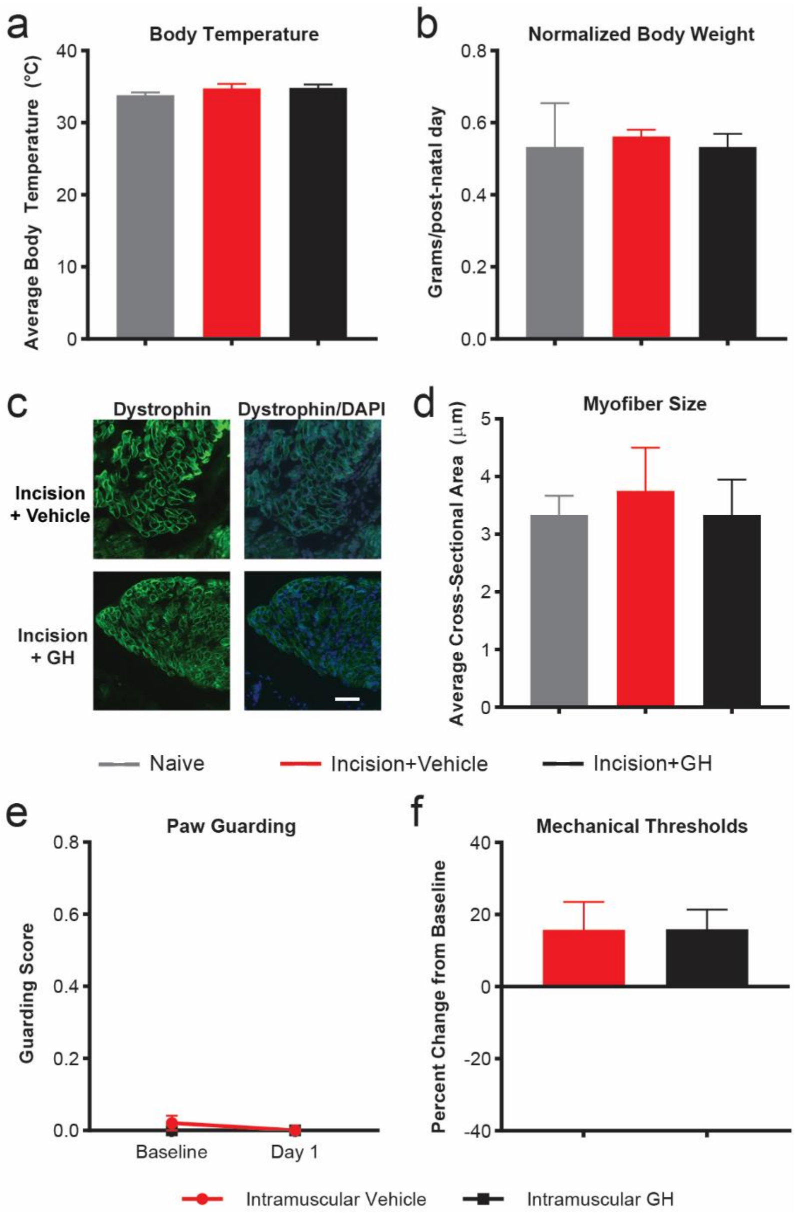


**Extended Figure 2-3: Off-target analyses of local GH injections.** a, Average body temperature measured on the thorax of neonates was not affected by injury or GH treatment. b, Body weight normalized to postnatal age was not affected by injury or GH injection. n = 4-6/group, one-way ANOVA. c, Representative images of myofibers stained with dystrophin (green) in both injured groups. DAPI (blue) co-stain was used to mark nuclei. d, The cross sectional area of injured animals was not affected by GH treatment. n = 3 animals/group, one-way ANOVA. e, Spontaneous paw guarding at Day 1 is not affected by local GH (1.5mg/kg)

injection in mice without incision compared to baseline (BL). f, Evoked mechanical withdrawal thresholds are not affected by local GH injection alone. n = 7-8/group, one-way ANOVA.

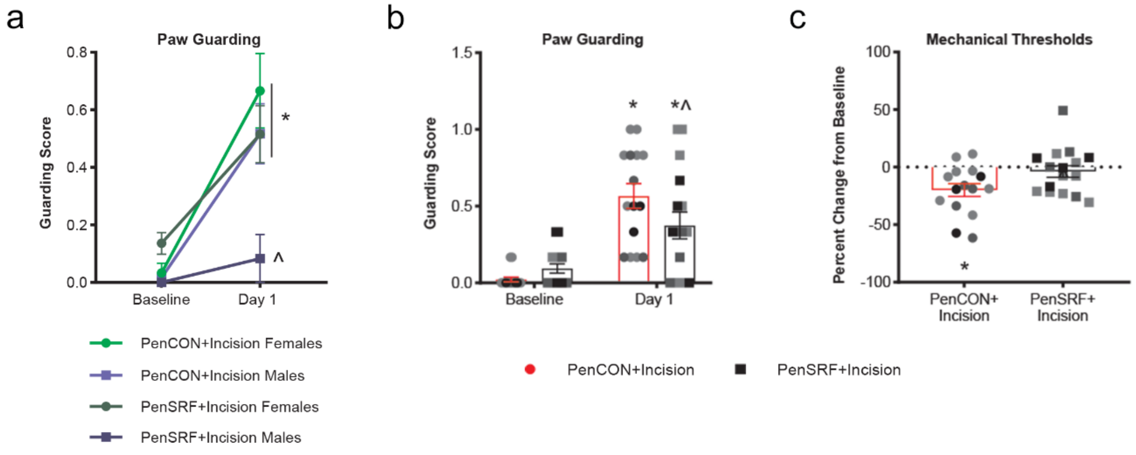


**Extended Figure 3-1: Sex-related effects of SRF inhibition on paw guarding after incision and age distribution for behavioral data in Figure 3**. a, Neonates injected with PenCON prior to an injury display increased guarding 1d after an incision. Females injected with PenSRF also have significantly increased guarding at 1d. Males injected with PenSRF do not differ from BL and guard significantly less than females. *p<0.001 vs. BL; ^p<0.001 vs. females and other Day 1 conditions. PenCON males = 10, females = 5; PenSRF males = 5, females = 11. Three-way RM ANOVA (GraphPad v.8) (time x sex x condition), Tukey's post hoc. Data shown as the mean ± s.e.m. b-c, Statistics and data points are equivalent to those displayed in Figure 3 where red bars indicates control group (circles) and black bars indicate experimental group (squares). Here, the shades of the data points indicate the age range of the animal at baseline behavior. Light gray = <u><</u>P7. Medium gray = P8 and P9. Dark gray = P10, P11. Black = <u>></u>P12.

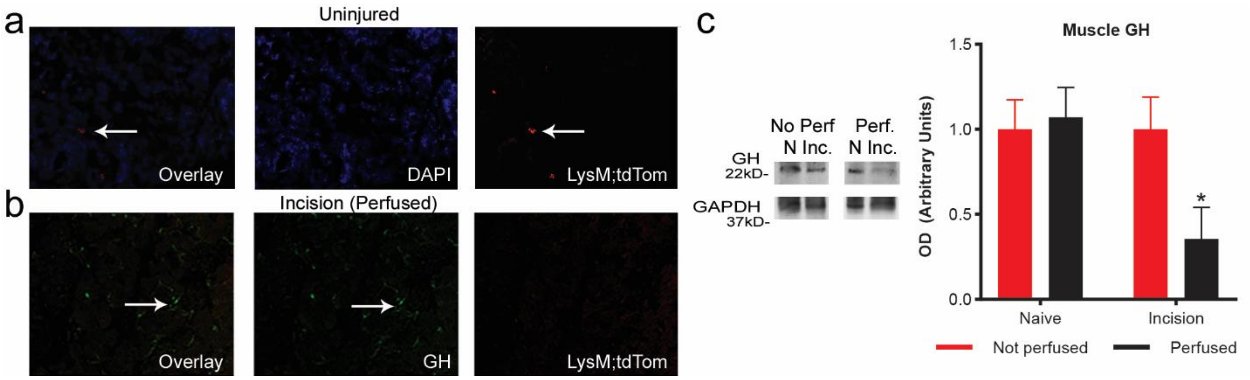


**Extended Figure 4-1: Cardiac perfusion with ice-cold saline reduces macrophages and consequently GH levels after a neonatal incision**. a, Representative images of uninjured hindpaw muscle from the macrophage reporter (LysM;tdTom) mouse co-stained with DAPI (blue) to mark nuclei. Arrows show few macrophages present in uninjured tissue. b, Images of injured hindpaw muscle in perfused LysM;tdTom animals (red) and stained for GH (green). Arrows indicate some remaining GH staining (compare to Figs. 4c, d). c, Representative western blots for GH in naïve and injured muscle with and without cardiac perfusion. Quantification of western blot analysis indicates that incised and perfused muscle have reduced GH levels compared to naïve unperfused. *$p<0.05$ vs. naïve unperfused. n = 4/group, two-way ANOVA, Tukey’s post hoc. Data shown as the mean ± s.e.m.

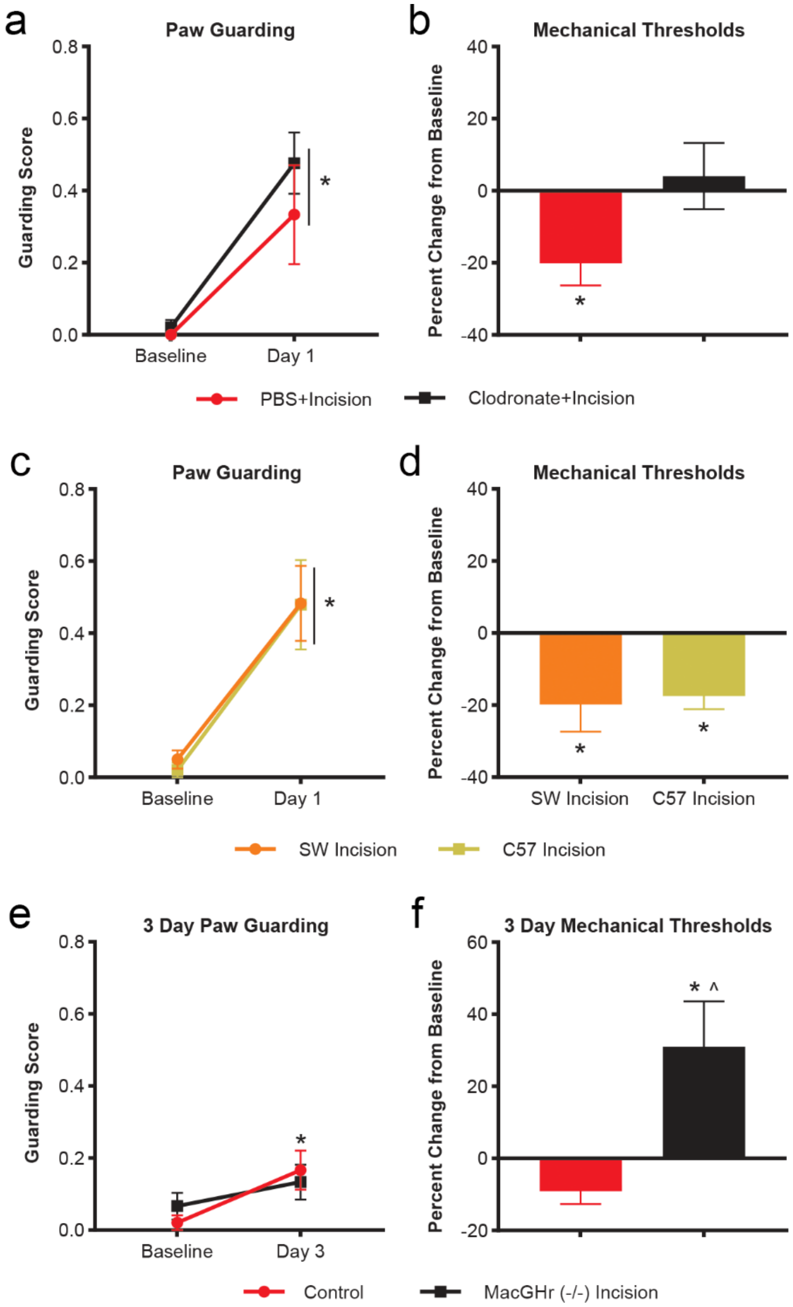


**Extended Figure 4-2: Behavioral effects of depleting macrophages by liposome encapsulated clodronate in neonates with hindpaw incision**. a, Animals guard after an injury regardless of intramuscular control PBS or clodronate injections. b, One day after an injury, PBS treated animals display reduced mechanical withdrawal thresholds, while clodronate treated animals do not. *p<0.05 vs. BL. n = 8/group, two-way RM ANOVA, Tukey's post hoc. c, One day after an injury, neonatal SW and C57 animals display spontaneous paw guarding compared to baseline values. d, Following an incision, neonatal animals from both strains display reduced squeezing withdrawal thresholds compared to baseline. *P<0.05 vs. BL. n = 8-10/group, two-way RM ANOVA, Tukey's post hoc.  e, Three days after a neonatal incision MacGHr-/- animals no longer display a difference from baseline (BL), but C57 animals still slightly guard their paw. f, Mechanical withdrawal thresholds are increased in MacGHr-/- 3 days after a neonatal injury compared to their BL and controls. *p<0.05 vs. BL; ^p<0.05 vs. controls. n = 8-10/group, two-way RM ANOVA, Tukey's post hoc. Data shown as the mean ± s.e.m.

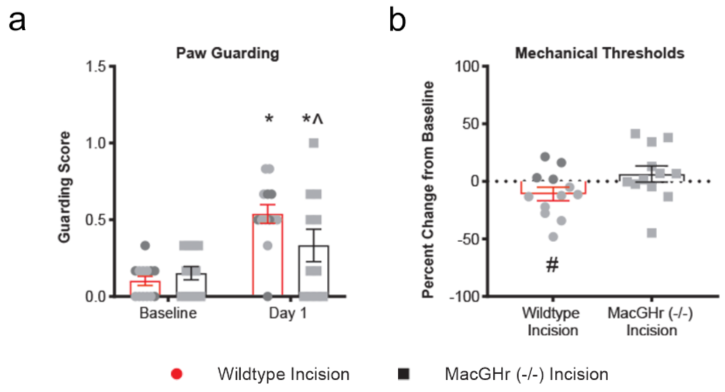


**Extended Data Figure 4-3: Age distribution for behavioral data in Figure 4.** Statistics and data points are equivalent to those displayed in Figure 4 where red bars indicates control group (circles) and black bars indicate experimental group (squares). Here, the shades of the data points indicate the age range of the animal at baseline behavior. Light gray = <u><</u>P7. Medium gray = P8 and P9. Dark gray = P10, P11. Black = <u>></u>P12.

**mRNA Primers**

| Gene | Forward | Reverse |
|---|---|---|
| ASIC3 | ATGAAACCTCCCTCAGGACTGG | AACTCCCCATAGTAGCGAACCC |
| ELK1 | AAGAATTGGAAGCTGCAAGGGCTG | TGTTCTCTGTTAGGATGGCTGGGA |
| ELK3 | ACACACACACAACCAAGATG | TCAGAGCGCTGGGATTATAG |
| ELK4 | GGTGATTTGTCGGAGAGTAG | CACCTCCTATCTCTGGGTAT |
| Fcer2a | GATCTAAGGAACGCCCAATC | CTGTGCGCTTCTCATTCA |
| GAPDH | ATGTGTCCGTCGTGGATCTGA | ATGCCTGCTTCACCACCTTCTT |
| GDNF | AGCTGCCAGCCCAGAGAATT | GCACCCCCGATTTTTGC |
| GHr | GCCTCTACACCGATGAGTAA | GGAAAGGACTACACCACCT |
| IGFr1 | TTGAACTTATGCGCATGTGCTGGC | TCTCATCCTTGATGCTGCCGATGA |
| IL1-r | AGGAATGTGGCTGAAGAGCACAGA | ACTCGTGTGACCGGATATTGCTTC |
| IL1β | TACAAGGAGAACCAAGCAAC | GGTGTGCCGTCTTTCATTA |
| MCP1 | CACCTGCTGCTACTCATTC | CTACAGCTTCTTTGGGACAC |
| NFκB | CTGCACCAAGACGGAACC | GAGCCTTCTCAAGAAAGAGGTTATC |
| NGF | ACACTCTGATCACTGCGTTTTTG | CCTTCTGGGACATTGCTATCTGT |
| OSMr | TCCAGGCTCACCCTTATT | AGCCTCGGTGTGTAGTT |
| P2X3 | ACAAGATGGAGAATGGCAGCGAGT | TGATGTTGAACTTGCCAGCGTTCC |
| P2Y1 | GATGAATTTGCGAGCACGGTTGGA | TCCACACAGCTGTTGAGACTTGCT |
| SRF | TGGAGTTCATCGACAACAAG | AGCGTGGACAGCTCATA |
| STAT1 | CCCAGGAATCTCTCCTTCTT | GACCTCTCTTGGTGACTGAT |
| STAT3 | CTGGGTCTGGCTAGACAATA | CGCTCCTTGCTGATGAAA |
| STAT5 | CCCACGTCAGTTGTAGTATC | GTTCAGCTCTTACACGAGAG |
| TNFa-r | TCGGAAAGAAATGTCCCAGGTGGA | TGGAACTGGTTCTCCTTACAGCCA |
| TNFα | CCTATGTCTCAGCCTCTTCT | GGGAACTTCTCATCCCTTTG |
| TRPA1 | GCAGGTGGAACTTCATACCAACT | CACTTTGCGTAAGTACCAGAGTGG |
| TRPV1 | TTCCTGCAGAAGAGCAAGAAGC | CCCATTGTGCAGATTGAGCAT |

**Extended Table 1-1: Primer information for standard realtime PCR.**

| Group III/IV Instantaneous Frequencies | | | | | | | | | |
|---|---|---|---|---|---|---|---|---|---|
| Afferent Type | Naïve | | | Incision + Vehicle | | | Incision + GH | | |
| (Hz) | Mean | s.e.m. | n | Mean | s.e.m. | n | Mean | s.e.m. | n |
| Mechanical | 73 | 13 | 21 | 111 | 27 | 13 | 59 | 12 | 22 |
| Cold | 18.8 | 10.4 | 9.0 | 44.1 | 13.6 | 10.0 | 29.8 | 9.3 | 10.0 |
| Hot | 59.1 | 1.8 | 2.0 | 51.3 | 21.6 | 3.0 | 78.2 | 39.7 | 8.0 |
| Low metabolite | 24.7 | 22.3 | 4.0 | 4.5 | n/a | 1.0 | 192.9 | 104.7 | 3.0 |
| High metabolite | 14.8 | 7.3 | 5.0 | 18.9 | 11.1 | 4.0 | 27.1 | 2.8 | 2.0 |
| Both metabolite | 7.8 | 4.0 | 4.0 | 43.2 | 20.6 | 6.0 | 0.7 | n/a | 1.0 |

| Group III/IV Thermal and Chemical Firing Rates | | | | | | | | | |
|---|---|---|---|---|---|---|---|---|---|
| Afferent Type | Naïve | | | Incision + Vehicle | | | Incision + GH | | |
| (Hz) | Mean | s.e.m. | n | Mean | s.e.m. | n | Mean | s.e.m. | n |
| Cold | 2.9 | 0.6 | 8.0 | 4.8 | 1.4 | 10.0 | 5.3 | 1.4 | 9.0 |
| Hot | 16.0 | n/a | 1.0 | 6.0 | 2.0 | 3.0 | 6.8 | 3.8 | 8.0 |
| Low metabolite | 2.4 | 0.9 | 7.0 | 2.1 | 0.5 | 7.0 | 2.8 | 1.0 | 5.0 |
| High metabolite | 3.4 | 0.1 | 7.0 | 5.8 | 3.0 | 9.0 | 1.0 | n/a | 1.0 |
| Both metabolite | 5.3 | 1.8 | 3.0 | 3.0 | 0.4 | 7.0 | 1.0 | n/a | 1.0 |

**Extended Table 2-1: Group III/IV instantaneous frequencies are unaffected between conditions within afferent subtypes.** One-way ANOVA with Tukey's or ANOVA on Ranks with Dunn's post hoc.
